# Conserved and divergent programs govern cerebellar output neuron development in zebrafish

**DOI:** 10.64898/2026.09.15.751670

**Authors:** Yukimi Koyama-Fujii, Chelsea U. Kidwell, Noga Moshe, Ivan Bassi, Karina Yaniv, Cecilia B. Moens, Takashi Shimizu, Masahiko Hibi

**Author notes:** Tune Therapeutics, 1930 Boren Ave., Suite 500, Seattle, WA 98101, USA.

## Abstract

Cerebellar output neurons transmit the results of cerebellar computation to other brain regions. In most vertebrates, cerebellar output neurons form discrete nuclei, whereas in teleosts they are represented by eurydendroid cells, which differ markedly in morphology and anatomical organization. How these divergent output systems are developmentally related remains unclear. Here, we show that zebrafish eurydendroid cells arise from *atoh1a*-expressing progenitors, including a population expressing *olig2*, revealing a developmental origin shared with cerebellar output neurons in other vertebrates. *atoh1a* is required for eurydendroid cell differentiation and regulates *lhx9* and *lhx2b*, which promote *robo1* and *robo3* expression and proper axon guidance. Our findings also suggest a role for Reelin signaling in eurydendroid cell positioning. Together, these findings reveal conserved developmental features shared by cerebellar output neurons across vertebrates, as well as mechanisms underlying the distinct anatomical organization of teleost eurydendroid cells.

## INTRODUCTION

The organization of the vertebrate brain has diversified throughout evolution ^1,2^. The cerebellum is no exception, exhibiting remarkable morphological and circuit diversity across vertebrate species while retaining a broadly conserved cellular organization ^3–5^. Purkinje cells and granule cells, the two principal neuronal populations of the cerebellum, are conserved throughout jawed vertebrates. Purkinje cells integrate inputs conveyed by granule cells and climbing fibers and transmit the resulting signals to cerebellar output neurons, which project to multiple brain regions to convey the computational output of the cerebellum.

Across vertebrates, cerebellar output neurons exhibit striking differences in their anatomical organization (Figure 1A). In mammals, cerebellar output neurons are grouped into paired deep cerebellar nuclei (DCN), whereas avian species possess both cerebellar nuclei and additional extracerebellar output nuclei ^6,7^. In contrast, in teleosts including zebrafish, cerebellar output neurons are distributed within the cerebellum, with their cell bodies located adjacent to Purkinje cells rather than forming discrete nuclei. These neurons extend elaborate dendrites into the molecular layer and have therefore been termed eurydendroid cells ^8–10^. Like the excitatory projection neurons of the mammalian DCN, teleost cerebellar output neurons are glutamatergic, receive inhibitory inputs from Purkinje cells, and project contralaterally to the red nucleus homolog, thalamus, and other targets ^10–13^. These shared anatomical and functional characteristics support the homology between teleost eurydendroid cells and the excitatory projection neurons of the amniote cerebellar nuclei despite their markedly different organization.

**Figure 1.**
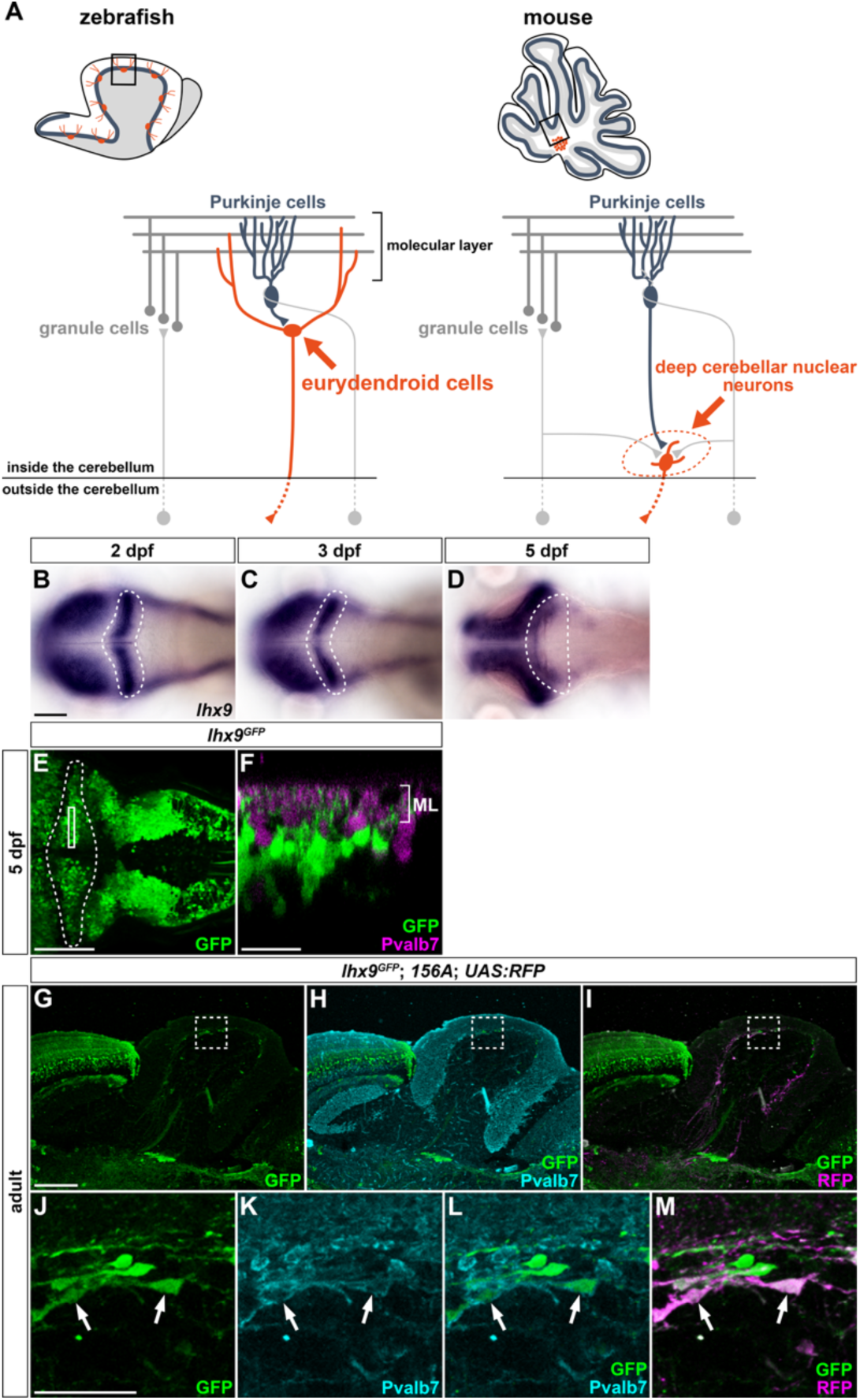
Expression of *lhx9* in eurydendroid cells. (A) Schematic of cerebellar organization and output neurons in zebrafish and mice. (B–D) Expression of *lhx9* in the cerebellum at 2 (B), 3 (C), and 5 dpf (D). In situ hybridization. Dorsal views with anterior to the left. The cerebellar region in each panel is outlined by a dashed line. (E, F) Expression of GFP (green; E, F) and parvalbumin 7 (Pvalb7; magenta; F) in 5-dpf *lhx9^GFP^* larvae. Immunostaining. Dorsal view (E) and z-sectional view (F). The cerebellar region in (E) is outlined by a dashed line. The rectangular box in (E) indicates the position of the z-section shown in (F). (G–M) Expression of GFP (green), Pvalb7 (cyan), and RFP (magenta) in the cerebellum of adult *lhx9^GFP^*; *hspzGFFgDMC156A*; *Tg(UAS:RFP)* zebrafish. Immunostaining of sagittal sections. (J–M) Higher-magnification views of the boxed regions in (G–I). Arrows indicate GFP- and RFP-positive eurydendroid cells surrounded by Pvalb7-positive Purkinje cell axons. ML, molecular layer. Scale bars: 100 μm (B, applies to B–D); 50 μm (E); 20 μm (F); 200 μm (G, applies to G–I); 50 μm (J, applies to J–M).

Whether these anatomical differences arose through modification of a conserved developmental program or through distinct developmental mechanisms remains unclear. In mammals, excitatory DCN neurons arise from progenitors expressing the proneural basic helix-loop-helix (bHLH) transcription factor Atoh1 in the upper rhombic lip (URL) and migrate to form the cerebellar nuclei ^14–17^. In contrast, many zebrafish cerebellar output neurons express *olig2*, which encodes a bHLH transcription factor ^18–20^. In both zebrafish and mice, *olig2* and *Olig2*, respectively, are expressed in ventricular zone (VZ) progenitors that generate Purkinje cells ^19,21^, leading to the hypothesis that at least a subset of teleost cerebellar output neurons originates from the VZ rather than the URL ^22^. However, direct evidence for the developmental origin of zebrafish cerebellar output neurons is lacking.

The distinctive organization of teleost cerebellar output neurons raises additional questions about the mechanisms that establish their position and axonal projections. In zebrafish, Reelin secreted by granule cells accumulates in the superficial molecular layer and regulates cerebellar neuronal migration. Mutations in *reelin* disrupt the positioning of both Purkinje cells and cerebellar output neurons ^23^. Cerebellar output neurons must also extend axons across the midline and toward distant brain targets, yet the molecular mechanisms that guide these projections remain poorly understood. Thus, whether the positioning and axon guidance of teleost cerebellar output neurons employ mechanisms shared with other neuronal populations or mechanisms associated with their distinctive organization remains unresolved.

Progress in addressing these questions has been hindered by the limited availability of molecular markers specific for zebrafish cerebellar output neurons. Although enhancer-trap screening identified a transgenic line labeling these neurons and enabled transcriptomic analyses ^10,24^, few molecular markers beyond *olig2* have been established. Recent single-cell transcriptomic analyses have begun to define the molecular identities of mammalian cerebellar nuclear neurons ^25,26^, but whether teleost cerebellar output neurons share transcriptional programs with their counterparts in other vertebrates remains unclear.

Here, we identify transcription factors that serve as markers for zebrafish eurydendroid cells and use these markers to define their developmental origin and the mechanisms controlling their differentiation, positioning, and axon guidance. We show that eurydendroid cells arise from *atoh1a*-expressing progenitors and uncover transcriptional mechanisms that regulate their differentiation and axon guidance, while Reelin signaling contributes to their positioning. Together, our findings reveal that, despite their divergent anatomy, teleost eurydendroid cells retain key developmental features of cerebellar output neurons in other vertebrates while employing mechanisms that contribute to their distinctive organization.

## RESULTS

### Eurydendroid cells express conserved cerebellar output neuron markers

We previously isolated the enhancer-trap line *hspzGFFgDMC156A* (hereafter referred to as *156A*), which expresses a modified version of GAL4 (GAL4FF) specifically in eurydendroid cells within the cerebellum ^10^. By crossing *156A* with *Tg(UAS:GFP)* (hereafter *UAS:GFP*), we labeled eurydendroid cells with GFP, isolated a cell population enriched for eurydendroid cells by fluorescence-activated cell sorting (FACS), and performed RNA sequencing (RNA-seq) ^24^. This analysis identified transcription factor genes enriched in the eurydendroid cell population compared with Purkinje cell and granule cell populations. Among these, *lhx9*, *meis2a*, and *meis2b* were of particular interest because their homologs are also expressed in excitatory cerebellar output neurons in mammals and birds (Table S1) ^25,26^.

Lhx9 is a transcription factor containing two LIM domains and a homeodomain ^27,28^. *lhx9* expression was detected in the superficial cerebellum at 2 days post-fertilization (dpf) and became progressively restricted to the anterior cerebellum by 5 dpf (Figures 1B– 1D). To label *lhx9*-expressing cells, we generated a knock-in line in which the *hsp70l* promoter-driven GFP cassette was inserted upstream of the *lhx9* transcription start site (*lhx9^GFP^*). In *lhx9^GFP^* larvae, GFP-positive cells did not express parvalbumin 7 (Pvalb7), a marker of Purkinje cells ^8^, but were located near Pvalb7-positive Purkinje cell bodies and extended dendrites into the molecular layer (Figures 1E and 1F; Figure S1). In adults, GFP-positive cells were similarly located near the Purkinje cell layer and were surrounded by axons of Pvalb7-positive Purkinje cells (Figures 1G–1M). Furthermore, GFP expression largely overlapped with *156A*-dependent RFP labeling (Figures 1G–1M), although a subset of GFP-positive cells lacked detectable RFP in both larvae and adults (Figure 1M; Figure S1), likely reflecting mosaic expression of the GAL4-UAS system ^29^. These observations establish *lhx9* as an early molecular marker of eurydendroid cells. The closely related Lhx2/9 family gene *lhx2b* showed a similar early developmental expression pattern, with expression in the superficial cerebellum at 2 dpf becoming progressively restricted to the anterior cerebellum by 5 dpf (Figure S1).

Meis2 is a member of the TALE homeodomain family of transcription factors ^30^. In zebrafish, it is encoded by two paralogous genes, *meis2a* and *meis2b*. During larval development, *meis2b* was prominently expressed in the cerebellum (Figures 2A–2C), while *meis2a* was also expressed in the larval cerebellum (Figure S2). Like *lhx9*, both *meis2* paralogs were expressed in the superficial cerebellum at 2 dpf and became restricted to the anterior cerebellum by 5 dpf. In adults, *meis2a* expression was detected near the Purkinje cell layer (Figure 2D). To determine whether these cells were eurydendroid cells, we generated *meis2b^GAL4^* and *meis2a^GFP^*knock-in reporter lines. In *meis2b^GAL4^*fish, GAL4-dependent GFP-labeled cells were located adjacent to Purkinje cells and extended dendrites into the molecular layer (Figures 2E and 2F; Figure S2). Similarly, in *meis2a^GFP^*fish, GFP-positive cells were located near Purkinje cells in both larvae and adults and were surrounded by axons of Pvalb7-positive Purkinje cells (Figures 2G–2M; Figure S2). All GFP-positive cells examined were also labeled by *156A*-dependent RFP (Figures 2I and 2M; Figure S2), confirming their identity as eurydendroid cells. Immunostaining with an antibody recognizing Meis2 revealed nuclear Meis2 immunoreactivity in cerebellar cells (Figures 2N– 2Q). Meis2-positive cells did not express Pvalb7, and their cell bodies were surrounded by Pvalb7-positive Purkinje cell axons (Figures 2N–2Q). Moreover, Meis2 immunoreactivity completely overlapped with GFP-positive cells in the cerebellum of *lhx9^GFP^* larvae (Figures 2R–2W), supporting the coexpression of *lhx9* and Meis2 in the same eurydendroid cell population. Together, these results establish *lhx9*, *meis2a*, and *meis2b* as molecular markers of zebrafish eurydendroid cells, with their homologs expressed in cerebellar output neurons in other vertebrates.

**Figure 2.**
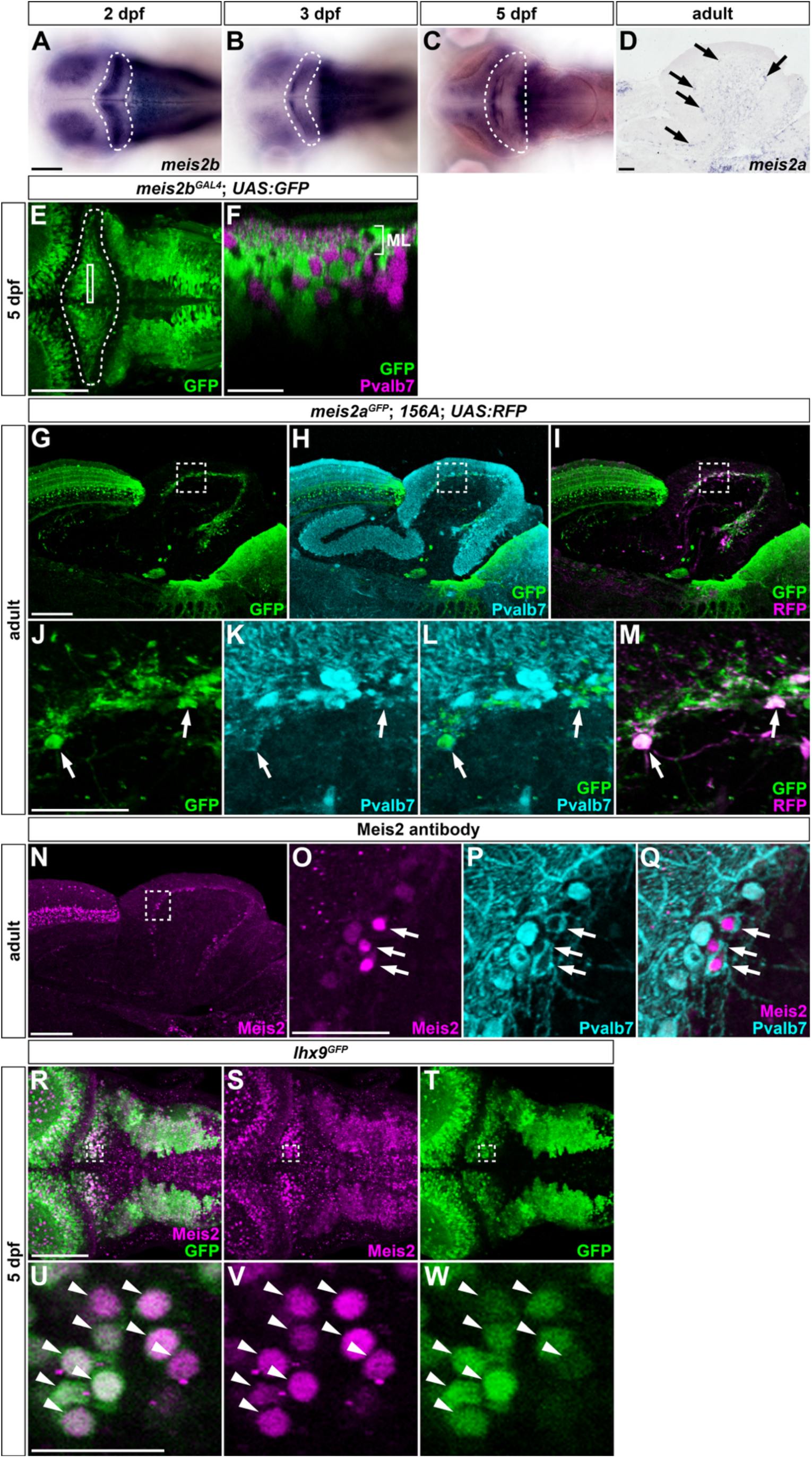
Expression of *meis2a* and *meis2b* in eurydendroid cells. (A–D) Expression of *meis2b* in the cerebellum at 2 (A), 3 (B), and 5 dpf (C), and expression of *meis2a* in the adult cerebellum (D). In situ hybridization. Dorsal views (A–C) and sagittal section (D). The cerebellar region in each of (A–C) is outlined by a dashed line. The position of *meis2a*-expressing eurydendroid cells is indicated by arrows in (D). (E, F) Expression of GFP (green; E, F) and parvalbumin 7 (Pvalb7; magenta; F) in a 5-dpf *meis2b^GAL4^*; *Tg(UAS:GFP)* larva. Immunostaining. Dorsal view (E) and z-sectional view (F). The cerebellar region in (E) is outlined by a dashed line. The rectangular box in (E) indicates the position of the z-section shown in (F). (G–M) Expression of GFP (green), Pvalb7 (cyan), and RFP (magenta) in the cerebellum of adult *meis2a^GFP^*; *hspzGFFgDMC156A*; *Tg(UAS:RFP)* zebrafish. Immunostaining of sagittal sections. (J–M) Higher-magnification views of the boxed regions in (G–I). Arrows indicate GFP- and RFP-positive eurydendroid cells surrounded by Pvalb7-positive Purkinje cell axons. (N–Q) Expression of Meis2 (magenta, N, O, Q) and Pvalb7 (cyan, P, Q) in the adult cerebellum. Immunostaining of sagittal sections. (O–Q) Higher-magnification views of the boxed region in (N). Arrows indicate Meis2-positive cells surrounded by Pvalb7-positive Purkinje cell axons. (R–W) Co-expression of Meis2 (magenta) and GFP (green) in the cerebellum of a 5-dpf *lhx9^GFP^* larva. Immunostaining. Dorsal views (R–T). (U–W) Higher-magnification views of the boxed regions in (R–T). Arrows indicate Meis2- and GFP-positive cells in the cerebellum. ML, molecular layer. Scale bars: 100 μm (A, applies to A– C); 100 μm (D); 50 μm (E); 20 μm (F); 200 μm (G, applies to G–I); 50 μm (J, applies to J– M); 200 μm (N); 50 μm (O, applies to O–Q); 50 μm (R, applies to R–T); 20 μm (U, applies to U–W).

### Eurydendroid cells arise from *atoh1*-expressing progenitors

In mammals, excitatory cerebellar output neurons arise from *Atoh1*-expressing progenitors in the upper rhombic lip (URL) ^14–17^, whereas many zebrafish eurydendroid cells express *olig2* ^18–20^. Zebrafish possess three *atoh1* genes, *atoh1a*, *atoh1b*, and *atoh1c*, which show partially overlapping but distinct expression patterns during cerebellar development ^19,31–33^. *atoh1c* is transiently expressed at the midbrain-hindbrain boundary (MHB) during the segmentation period (14–22 hpf) ^33^. From approximately 2 dpf, *atoh1a* and *atoh1b* are expressed in the URL. *atoh1a* expression subsequently becomes restricted to the anterior URL and, by 2.5–3 dpf, to the anterior cerebellum corresponding to the valvula cerebelli, where its expression persists into adulthood ^19,33^. *olig2* is also expressed in the cerebellar region from approximately 2 dpf ^19^. Given the prominent expression of *atoh1a* in the URL during the period of eurydendroid cell development, we examined its relationship with *olig2* expression. We first analyzed their expression patterns by in situ hybridization. To label *atoh1a*-expressing cells, we generated an *atoh1a^GAL4^* knock-in line in which the *hsp70l* promoter-driven GAL4FF cassette was inserted upstream of the *atoh1a* transcription start site. We then compared *atoh1a^GAL4^*-driven mCherry expression from *UAS:mCherry-CreERT2* with GFP expression from *TgBAC(olig2:EGFP)* (hereafter *olig2:GFP*).

Consistent with previous reports ^19,32,33^, *atoh1a* was broadly expressed in the URL at 2 dpf and became progressively restricted to the anterior cerebellum during subsequent development (Figures 3A–3C). In *atoh1a^GAL4^*; *UAS:mCherry-CreERT2* larvae, mCherry expression at 2 dpf recapitulated the broad URL expression of *atoh1a* mRNA (Figures 3G and 3J). At later stages, mCherry-positive cells remained distributed throughout the URL and were also found ventral to the URL, consistent with ventral migration (Figures 3H, 3I, 3K, and 3L). The persistence of mCherry labeling presumably reflects the stability of mCherry in cells with previous *atoh1a*-driven GAL4 activity. At 2 dpf, *olig2* was expressed not only in the posterior ventricular zone (VZ) but also in the posterior URL (Figures 3D, 3Da, and 3Db), with cerebellar expression persisting during subsequent larval development (Figures 3E and 3F). In *olig2:GFP* larvae, GFP-positive cells were subsequently detected slightly ventral to the URL, at the position occupied by eurydendroid cells (Figures 3M–3R). Notably, these *olig2:GFP*-positive cells were also labeled with mCherry in *atoh1a^GAL4^*; *UAS:mCherry-CreERT2* larvae (Figures 3S–3X). These observations raised the possibility that *olig2*-expressing eurydendroid cells arise from progenitors that express both *atoh1a* and *olig2* (Figure 3Y).

**Figure 3.**
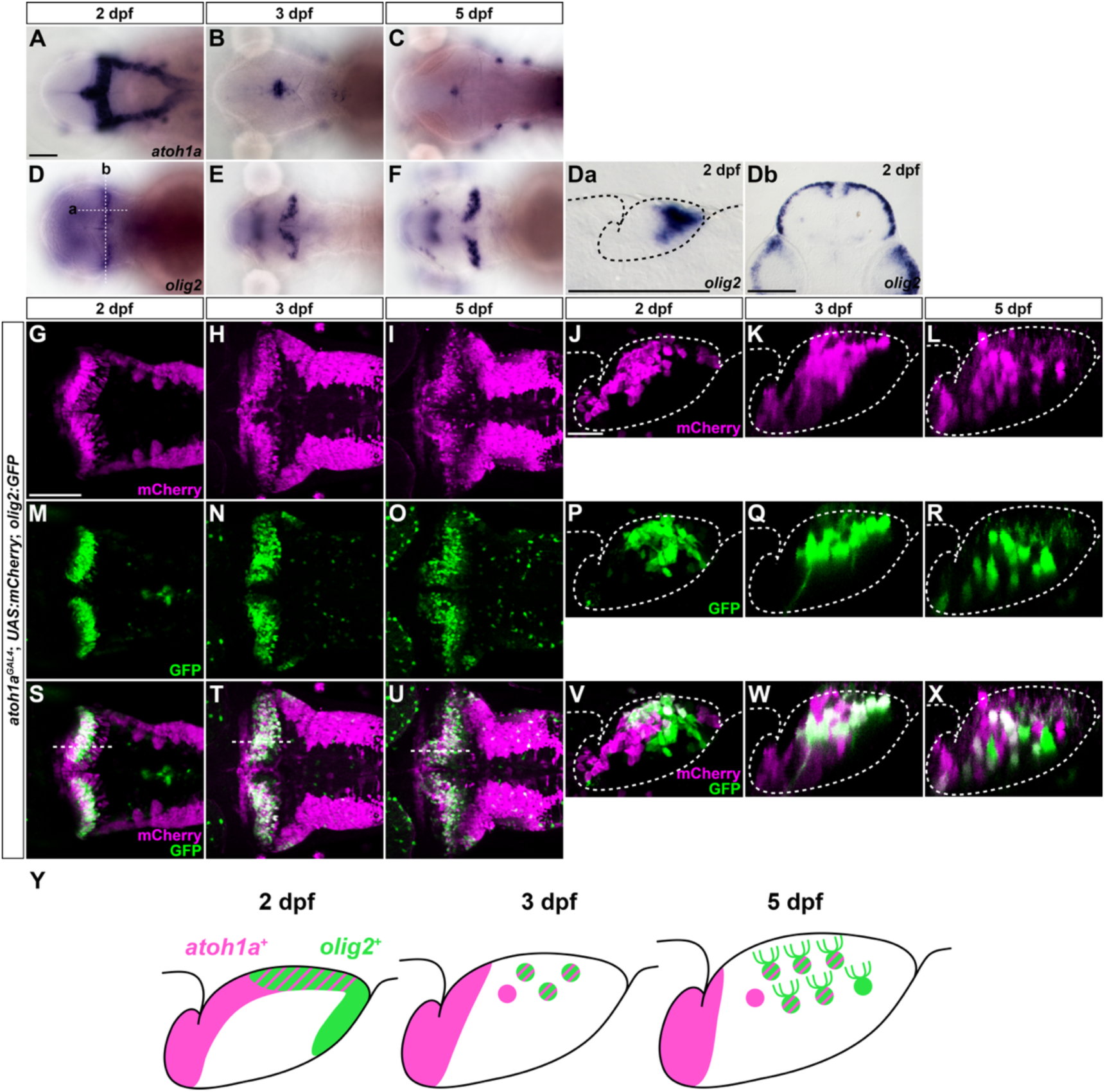
Expression of *atoh1a* and *olig2* during eurydendroid cell development. (A–F) Expression of *atoh1a* (A–C) and *olig2* (D–F) at 2 (A, D), 3 (B, E), and 5 dpf (C, F). In situ hybridization. Dorsal views. (Da, Db) Sagittal (Da) and transverse (Db) sections at the positions indicated in (D). (G–X) Expression of mCherry (magenta) and GFP (green) in 2-dpf (G, J, M, P, S, V), 3-dpf (H, K, N, Q, T, W), and 5-dpf (I, L, O, R, U, V) *atoh1a^GAL4^*; *Tg(UAS:mCherry-2A-CreERT2)*; *TgBAC*(*olig2:EGFP)* larvae. Immunostaining. Dorsal views (G–I, M–O, S–U), sagittal sections (J, P, V), and z-sectional views (K, L, Q, R, W, X). The approximate section planes at 2, 3, and 5 dpf are indicated in (S), (T), and (U), respectively. The 2-dpf dorsal and sagittal views were obtained from different larvae. Dashed outlines delineate the cerebellum in the section views. (Y) Schematic showing the localization of *atoh1a*- and *olig2*-expressing progenitors and differentiating cells derived from *atoh1a*- and/or *olig2*-expressing progenitors in the cerebellum at 2, 3, and 5 dpf. Scale bars: 100 μm (A, applies to A–F); 100 μm (Da); 100 μm (Db); 50 μm (G, applies to G–I, M–O, and S–U); 20 μm (J, applies to J–L, P–R, and V–X).

To directly determine whether eurydendroid cells arise from neural progenitors expressing *atoh1*, *olig2*, or *ptf1a*, we performed inducible genetic lineage tracing. In addition to the *atoh1a^GAL4^* line described above, we generated an *atoh1b^GAL4^* knock-in line using the same strategy and a *TgBAC(olig2:GAL4FF)* line (hereafter *olig2:GAL4*) for this analysis. We used these lines together with *TgBAC(atoh1c:GAL4FF)* (hereafter *atoh1c:GAL4*) ^33^ and *TgBAC(ptf1a:GAL4-VP16)* (hereafter *ptf1a:GAL4*) ^34^ to drive CreERT2 expression from *Tg(UAS:mCherry-CreERT2)* ^35,36^. To visualize recombined eurydendroid cells, which express *vglut2a*, we additionally introduced the *Tg(vglut2a:LOXP-DsRed-LOXP-GFP)* reporter ^37^. Endoxifen treatment induces CreERT2-mediated recombination in cells expressing each GAL4 driver, switching reporter expression from DsRed to GFP and thereby allowing *vglut2a*-expressing descendants of the recombined cells to be identified (Figure 4A).

**Figure 4.**
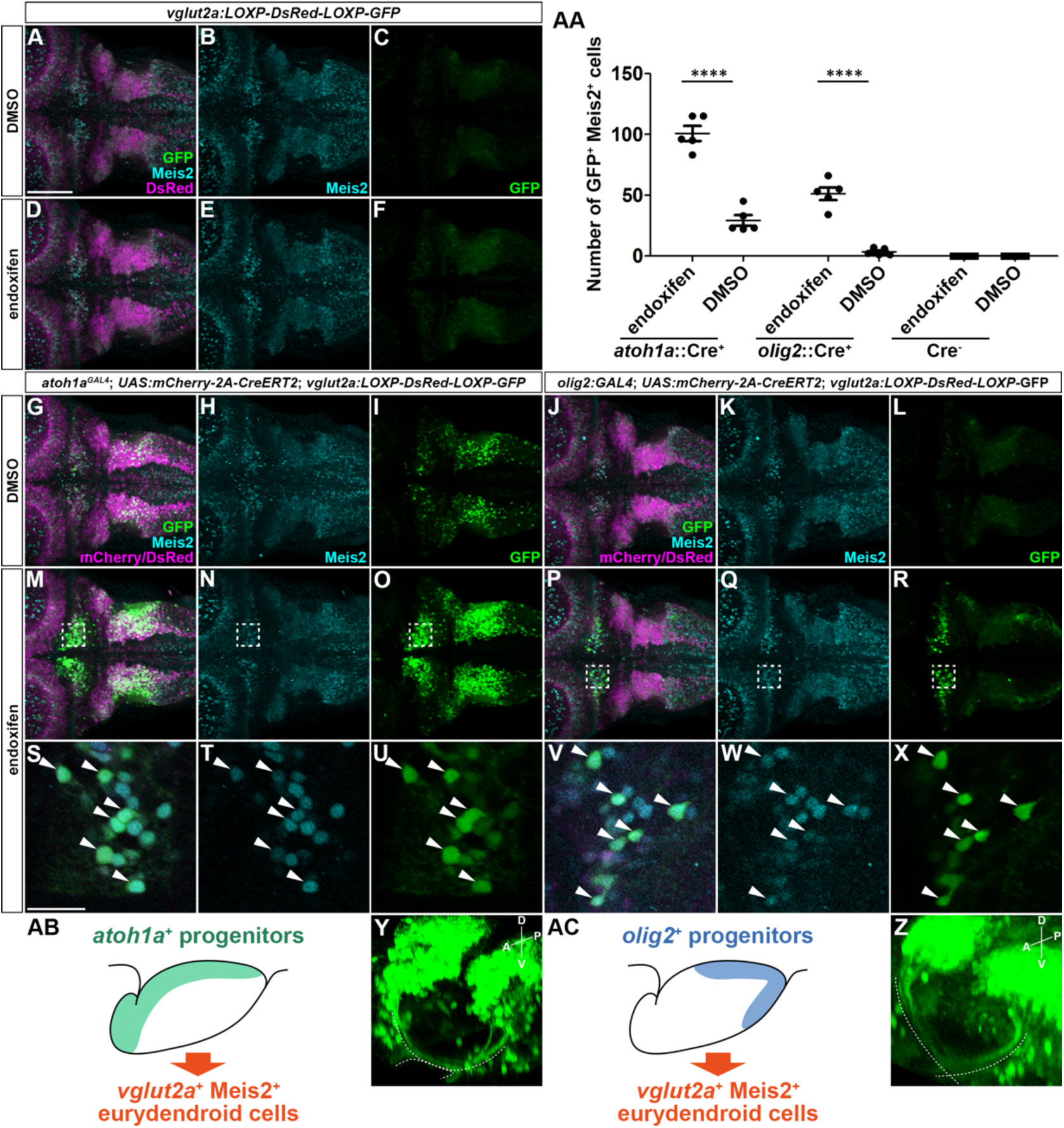
Eurydendroid cells are derived from *atoh1a*- and *olig2*-expressing cells. (A–F) Expression of GFP (green), Meis2 (cyan), and DsRed (magenta) in the cerebellar region of 5-dpf *Tg(vglut2a:LOXP-DsRed-LOXP-GFP)* (reporter Tg) larvae. Larvae were treated with DMSO (control) or endoxifen for 16 h from 2.5 dpf. Immunostaining for GFP and Meis2. Dorsal views. (G–Z) Lineage tracing in 5-dpf *atoh1a^GAL4^*; *Tg(UAS:mCherry-2A-CreERT2)*; reporter Tg larvae (G–I, M–O, S–U, Y) or *TgBAC(olig2:GAL4FF)*; *Tg(UAS:mCherry-2A-CreERT2)*; reporter Tg larvae (J–L, P–R, V–X, Z). Larvae were treated with DMSO or endoxifen for 16 h from 2.5 dpf. (G–X) Expression of GFP (green), Meis2 (cyan), and mCherry/DsRed (magenta). Immunostaining for GFP and Meis2. Dorsal views (G–X). (S–U) Higher-magnification views of regions corresponding to the boxed areas in (M–O), obtained from another larva. (V–X) Higher-magnification views of the boxed regions in (P–R). Arrowheads indicate GFP-positive, Meis2-positive cells. (Y, Z) GFP-labeled eurydendroid cells and their axonal projections following endoxifen treatment. Oblique dorsolateral views of the cerebellar region. White dotted lines indicate eurydendroid cell axon trajectories. A, anterior; P, posterior; D, dorsal; V, ventral. (AA) Numbers of *vglut2a*:GFP^+^ Meis2^+^ cells labeled by lineage tracing using the *atoh1a* and *olig2* driver lines and the control reporter line. Cre⁺ indicates larvae carrying both the GAL4 driver and *Tg(UAS:mCherry-2A-CreERT2)* transgenes, whereas Cre⁻ indicates larvae lacking one or both of these transgenes, as determined by genotyping. Data are shown as mean ± s.e.m., *n* = 5 larvae per condition. Each symbol represents one larva. Statistical analyses were performed separately for the *atoh1a* and *olig2* lineage-tracing experiments using two-way ANOVA with CreERT2 expression competence (Cre⁺ or Cre⁻) and treatment as factors, followed by Tukey’s multiple-comparison test. \*\*\*\**P* < 0.0001. (AB, AC) Schematics of lineage tracing of eurydendroid cells from *atoh1a*- and *olig2*-expressing progenitors. Scale bars: 50 μm (A, applies to A–R); 20 μm (S, applies to S–X).

We first induced recombination by treating larvae with endoxifen for 16 h from 2.5 dpf. In Cre⁻ control larvae, which lacked one or both transgenes required for CreERT2 expression, GFP-positive cells were rarely detected in the cerebellum (Figures 4A–4F and 4AA). In larvae expressing CreERT2 under the control of *atoh1a^GAL4,^* a small number of cerebellar GFP-positive cells were detected even in DMSO-treated controls, whereas endoxifen treatment resulted in numerous GFP-positive cells in the cerebellum. Nearly all of these GFP-positive cells expressed Meis2 (Figures 4G–4I, 4M–4O, 4S–4U, and 4AA), identifying them as eurydendroid cells. Similarly, lineage tracing with *olig2:GAL4* yielded few GFP-positive cells in the absence of endoxifen but numerous GFP-positive cells following endoxifen treatment (Figures 4J–4L, 4P–4R, 4V–4X, and 4AA). These cells were also Meis2 positive. Furthermore, GFP-positive cells labeled by either the *atoh1a* or *olig2* lineage extended axons anteroventrally from the cerebellum and crossed the midline, a characteristic projection pattern of eurydendroid cells (Figures 4Y and 4Z). Together with the overlapping *atoh1a* and *olig2* expression described above, these results indicate that eurydendroid cells arise from *atoh1a*-expressing progenitors and that this lineage includes cells that express *olig2*.

We next performed the same lineage-tracing experiments using *atoh1b^GAL4^*, *atoh1c:GAL4*, and *ptf1a:GAL4*. The *atoh1b* lineage also gave rise to GFP-positive, Meis2-positive cells, although fewer such cells were observed than with the *atoh1a* driver (Figure S3). With *atoh1c:GAL4*, few GFP-positive, Meis2-positive cells were detected following endoxifen treatment for 16 h from 2.5 dpf, whereas such cells were detected when recombination was induced earlier, from 12 to 36 hpf (Figure S4), consistent with the transient early expression of *atoh1c*. In contrast, lineage tracing with *ptf1a:GAL4* labeled very few GFP-positive, Meis2-positive cells per larva (Figure S5). The same *ptf1a*:*GAL4* line efficiently labels Purkinje cell lineages ^35^, demonstrating its activity in the cerebellum. Because labeling efficiency may nevertheless differ among GAL4 driver lines, the relative contributions of the lineages examined cannot be quantitatively compared. Despite this limitation, the extensive labeling of eurydendroid cells in *atoh1a* lineage tracing, together with the identification of eurydendroid cells derived from the *atoh1b* and early *atoh1c* lineages, establishes that eurydendroid cells arise from *atoh1*-expressing progenitors (Figure 4AB). A subset of eurydendroid cells also arises from *olig2*-expressing progenitors (Figure 4AC).

### Atoh1a is required for eurydendroid cell differentiation

To determine the roles of *atoh1* genes in eurydendroid cell differentiation, we analyzed zebrafish carrying mutations in each of the three *atoh1* paralogs. The previously described *atoh1a^fh282^* allele contains a point mutation resulting in an amino acid substitution within the basic helix-loop-helix (bHLH) domain ^33^. We therefore generated two additional *atoh1a* alleles, *atoh1a^Δ23^* and *atoh1a^Δ4^*, using CRISPR/Cas9. Both deletions cause frameshifts, and the resulting mutant alleles are predicted to encode truncated proteins lacking most of the bHLH domain. We also generated the *atoh1b^fh511^* and *atoh1b^Δ2^* alleles and used the previously described *atoh1c^fh367^* allele; all three are predicted to encode proteins lacking the bHLH domain (Figure S6). Expression of *atoh1a* and *atoh1c* was not detectably altered in *atoh1a^Δ23^* and *atoh1c^fh367^* mutants, respectively (Figure S7), suggesting that these mutant transcripts are not subject to substantial nonsense-mediated decay. To assess eurydendroid cell differentiation, we analyzed GFP expression at 5 dpf using *156A*; *UAS:GFP* or *lhx9^GFP^* in single and compound *atoh1* mutants (Figures 5A–5J), as well as Meis2 expression (Figures 5K–5O; Figure S8).

**Figure 5.**
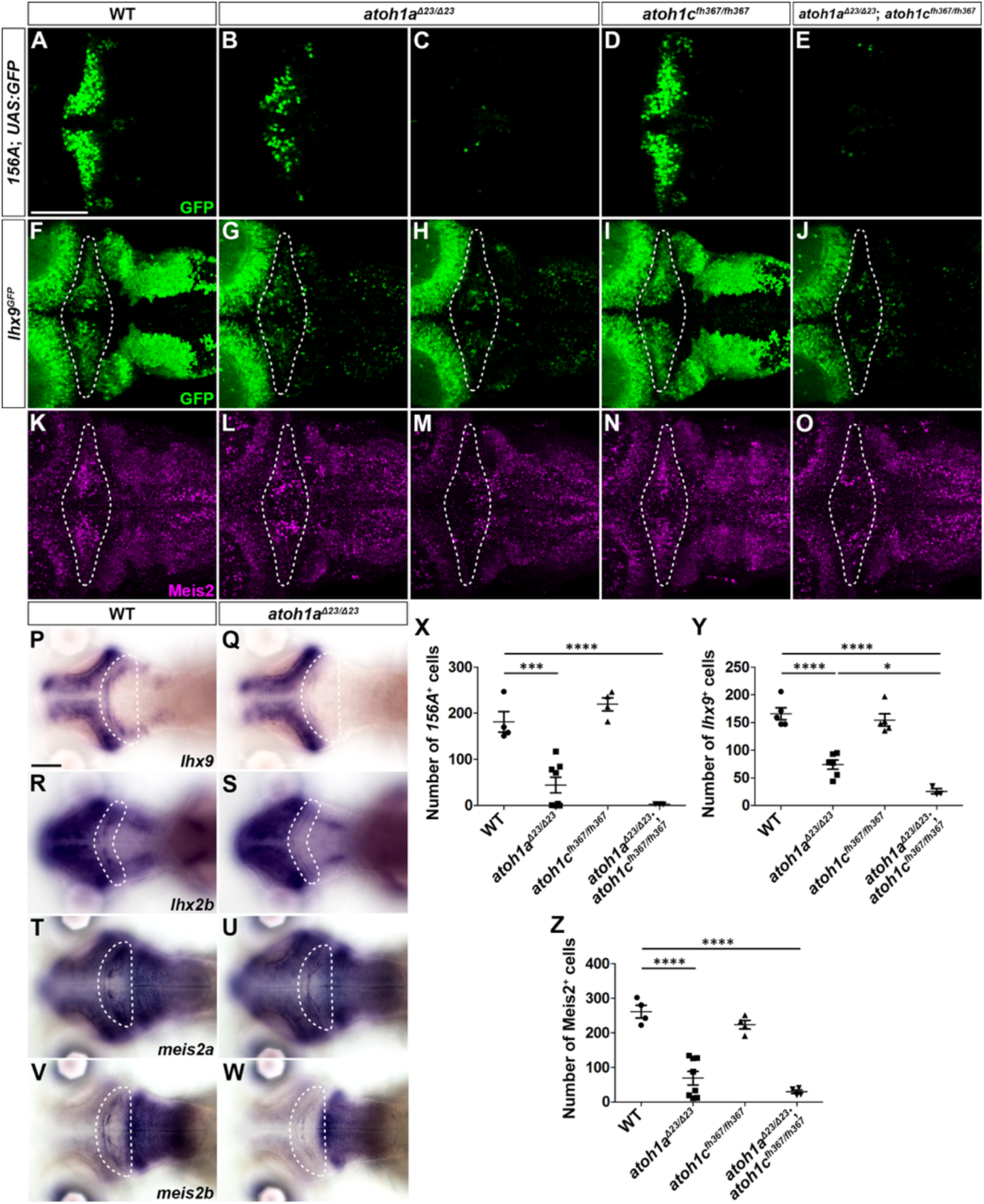
Atoh1a is required for differentiation of eurydendroid cells. (A–E) Expression of GFP (green) in 5-dpf wild-type (WT; A, *n* = 4), *atoh1a^Δ23^* (B, *n* = 4; C, *n* = 4), *atoh1c^fh367^* (D, *n* = 4), and *atoh1a^Δ23^*; *atoh1c^fh367^* (E, *n* = 4) *hspzGFFgDMC156A*; *Tg(UAS:GFP)* larvae. (F–J) Expression of GFP (green) in 5-dpf WT (F, *n* = 5), *atoh1a^Δ23^* (G, *n* = 4; H, *n* = 2), *atoh1c^fh367^* (I, *n* = 5), and *atoh1a^Δ23^*; *atoh1c^fh367^* (J, *n* = 3) *lhx9^GFP^* larvae. (K–O) Expression of Meis2 (magenta) in 5-dpf WT (K, *n* = 4), *atoh1a^Δ23^* (L, *n* = 4; M, *n* = 4), *atoh1c^fh367^* (N, *n* = 4), and *atoh1a^Δ23^*; *atoh1c^fh367^* (O, *n* = 4) larvae. All mutants were homozygous for the indicated alleles. Dorsal views. Immunostaining. The cerebellar region in F–O is outlined by a dashed line. Two phenotypic classes were observed in *atoh1a^Δ23^*mutant larvae: one showing a marked reduction in eurydendroid cell markers (B, G, L) and the other showing little or no marker expression in the cerebellum (C, H, M). (P–W) Expression of *lhx9* (P, Q), *meis2a* (T, U), and *meis2b* (V, W) in 5-dpf WT (P, *n* = 3; T, *n* = 6; V, *n* = 5) and *atoh1aΔ23* mutant larvae (Q, *n* = 6; U, *n* = 6; W, *n* = 5), and expression of *lhx2b* (R, S) in 3-dpf WT (R, *n* = 5) and *atoh1a^Δ23^* mutant larvae (S, *n* = 5). In situ hybridization. Dorsal views. The cerebellar region in each panel is outlined by a dashed line. (X) Numbers of GFP-positive cells in WT and mutant *hspzGFFgDMC156A*; *Tg(UAS:GFP)* larvae. (Y) Numbers of GFP-positive cells in WT and mutant *lhx9^GFP^* larvae. (Z) Numbers of Meis2-positive cells in the cerebellum of WT and mutant larvae. Data are shown as mean ± s.e.m. Each symbol represents one larva. \**P*< 0.05, \*\*\**P* < 0.001, \*\*\*\**P* < 0.0001 (One-way ANOVA with Tukey’s multiple-comparison test). Scale bars: 50 μm (A, applies to A–O); 100 μm (P, applies to P–W).

Compared with wild-type larvae, *atoh1a^Δ23^* homozygous mutants showed a marked reduction in GFP-positive eurydendroid cells labeled by either *156A*; *UAS:GFP* or *lhx9^GFP^*. Two phenotypic classes were observed. In one group, GFP-positive cells were almost completely absent (4/8 larvae for *156A*; *UAS:GFP* and 2/6 for *lhx9^GFP^*; Figures 5C and 5H), whereas the remaining mutants retained a small number of GFP-positive cells (4/8 and 4/6 larvae, respectively; Figures 5B and 5G). A similar variation was observed for Meis2-positive cells: these cells were almost completely absent in 4/8 mutants (Figure 5M), whereas appreciable numbers remained in the other 4/8 mutants (Figure 5L). A similarly variable reduction in Meis2-positive eurydendroid cells was observed in *atoh1a^Δ4^* homozygous mutants (Figure S8). Notably, the severe loss of Meis2-positive cells observed in some *atoh1a^Δ23^* and *atoh1a^Δ4^* mutants was not observed in *atoh1a^fh282^* mutants (Figure S8). By contrast, homozygous *atoh1b* or *atoh1c* single mutants showed no obvious reduction in eurydendroid cells (Figures 5D, 5I, 5N, and S8).

The variable phenotype of *atoh1a^Δ23^* mutants raised the possibility that other *atoh1* paralogs compensate for loss of *atoh1a*. Notably, a subset of *atoh1a^Δ23^* homozygous mutants showed elevated *atoh1c* expression (Figure S7). The number of Meis2-positive cells did not differ significantly between *atoh1a^Δ23^* single mutants and *atoh1a^Δ23^*; *atoh1b^Δ2^* double homozygous mutants (Figure S8). In striking contrast, all *atoh1a^Δ23^*; *atoh1c^fh367^* double homozygous mutants showed a nearly complete loss or a dramatic reduction of GFP-positive eurydendroid cells labeled by either *156A*; *UAS:GFP* or *lhx9^GFP^*, as well as Meis2-positive eurydendroid cells (Figures 5X–5Z). These results indicate that *atoh1a* plays the predominant role in eurydendroid cell differentiation and support a compensatory role for *atoh1c* in the absence of *atoh1a*.

We next examined whether the expression of transcription factors associated with eurydendroid cell differentiation depends on Atoh1a. In *atoh1a^Δ23^* homozygous mutants, cerebellar expression of *lhx2b*, *lhx9*, *meis2a*, and *meis2b* was markedly reduced compared with wild-type larvae (Figures 5P–5W). These findings suggest that these eurydendroid cell transcription factor genes are regulated downstream of Atoh1a.

### Lhx9 and Lhx2b control eurydendroid cell axon guidance

To investigate the roles of the eurydendroid cell transcription factors Lhx9 and Lhx2b, we generated mutant alleles of *lhx9* and *lhx2b* using CRISPR/Cas9. The *lhx9^Δ16^* allele is predicted to encode a truncated Lhx9 protein that terminates within the second LIM domain and lacks the homeodomain. Similarly, the *lhx2b^Δ2^*allele is predicted to encode a truncated Lhx2b protein lacking most of the first LIM domain, the second LIM domain, and the homeodomain (Figure S9). We introduced these mutations into fish carrying *156A*; *UAS:GFP* and examined the morphology and axonal projections of eurydendroid cells.

In wild-type larvae, GFP-positive eurydendroid cell axons crossed the midline and then extended anteriorly on the contralateral side toward the forebrain, with projections reaching the presumptive thalamic, preoptic, and hypothalamic regions (Figures 6A–6C, 6M, and 6O), consistent with cerebellar projection targets reported in goldfish ^38^. Eurydendroid cell axons from opposite sides crossed the midline in opposite directions (Figures 6A–6C and 6P). Neither *lhx9^Δ16^* nor *lhx2b^Δ2^* single homozygous mutants showed obvious abnormalities in these projections (Figures 6D–6I, 6O, and 6P). In contrast, *lhx9^Δ16^*; *lhx2b^Δ2^* double homozygous mutants showed a marked reduction in axon bundles crossing the midline, with many axons instead extending anteriorly on the ipsilateral side (Figures 6J–6L and 6P). Rostral projections were also markedly reduced in the double mutants, particularly those extending toward the presumptive preoptic and hypothalamic regions (Figures 6J, 6N, and 6O). These results indicate that Lhx9 and Lhx2b act redundantly to control midline crossing and the formation of rostral projections by eurydendroid cell axons.

**Figure 6.**
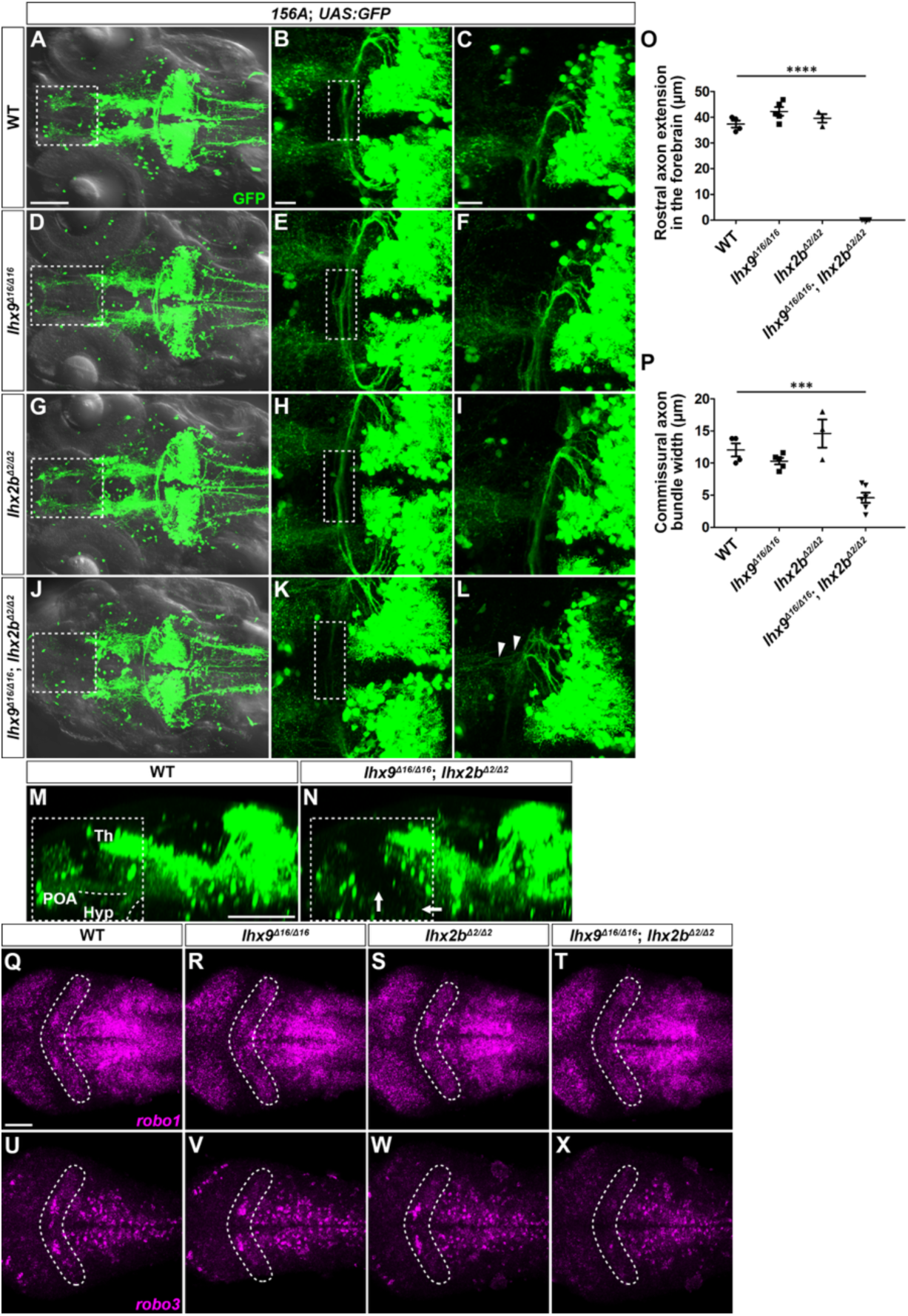
*lhx9* and *lhx2b* are required for proper axon guidance of eurydendroid cells. (A–L) Expression of GFP (green) in 5-dpf wild-type (WT; A–C, *n* = 4), *lhx9*^Δ16^ (D–F, *n* = 5), *lhx2b*^Δ2^ (G–I, *n* = 3), and *lhx9*^Δ16^; *lhx2b*^Δ2^ (J–L, *n* = 6) *hspzGFFgDMC156A*; *Tg(UAS:GFP)* larvae. All mutants were homozygous for the indicated alleles. Larvae were immunostained for GFP and cleared before imaging. Dorsal views of confocal z-stack projections. (A, D, G, J) Overview images of the forebrain, midbrain, and hindbrain. Dashed boxes indicate the regions containing axonal projections within the forebrain. (B, E, H, K) Higher-magnification views of the midbrain–cerebellum boundary region. Dashed boxes indicate the regions where axons from eurydendroid cells on the left and right sides cross the midline. (C, F, I, L) Higher-magnification views of the regions where eurydendroid cell axons exit the cerebellum. Arrowheads indicate axons that fail to cross the midline and instead project anteriorly on the ipsilateral side. (M, N) Lateral confocal images showing GFP-positive eurydendroid cell axons in the same WT and *lhx9*^Δ16^; *lhx2b*^Δ2^ double mutant larvae shown in (A) and (J), respectively. Dashed boxes indicate the regions containing axonal projections within the forebrain, corresponding to those marked in the dorsal views (A, J). Dashed lines in (M) indicate axonal projections toward the presumptive preoptic and hypothalamic regions. Arrows in (N) indicate the corresponding regions, where these projections are markedly reduced in the double mutant. Th, thalamus; POA, preoptic area; Hyp, hypothalamus. (O) Rostral extent of axonal projections within the forebrain, measured from the forebrain–midbrain boundary to the anterior ends of the axons. (P) Width of the axon bundle formed by eurydendroid cell axons crossing the midline. Data are shown as mean ± s.e.m. Each symbol represents one larva. \*\*\**P* < 0.001, \*\*\*\**P* < 0.0001 (one-way ANOVA with Tukey’s multiple-comparison test). (Q–X) Expression of *robo1* (Q–T) and *robo3* (U–X) (magenta) in 3-dpf larvae. Fluorescent in situ hybridization. Dorsal views, with rostral to the left. *robo1* expression is shown in WT (Q, *n* = 5), *lhx9^Δ16^* (R, *n* = 5), *lhx2b^Δ2^* (S, *n* = 5), and *lhx9*^Δ16^; *lhx2b*^Δ2^ (T, *n* = 5) larvae. *robo3* expression is shown in WT (U, *n* = 5), *lhx9*^Δ16^ (V, *n* = 5), *lhx2b^Δ2^* (W, *n* = 5), and *lhx9*^Δ16^; *lhx2b*^Δ2^ (X, *n* = 5) larvae. The cerebellar region in each of (Q–X) is outlined by a dashed line. The images shown are representative of the indicated numbers of larvae. Scale bars: 100 μm (A, applies to A, D, G, J); 20 μm (B, applies to B, E, H, K); 20 μm (C, applies to C, F, I, L); 100 μm (M, applies to M and N); 50 μm (Q, applies to Q–X).

Midline guidance of commissural axons is regulated by Slit-Robo signaling, and we previously found that *robo1* and *robo3* are expressed in the zebrafish cerebellum ^39^. We therefore examined whether their expression was affected by loss of Lhx9 and Lhx2b. Both *robo1* and *robo3* were expressed in the cerebellum of wild-type larvae, and their expression was not obviously altered in either *lhx9^Δ16^* or *lhx2b^Δ2^* single mutants. In *lhx9^Δ16^*; *lhx2b^Δ2^* double mutants, however, cerebellar expression of both *robo1* and *robo3* was markedly reduced (Figures 6Q–6X). Together, these findings identify Lhx9 and Lhx2b as redundant regulators of eurydendroid cell axon guidance and suggest that they act, at least in part, by promoting *robo1* and *robo3* expression.

### Reelin signaling contributes to eurydendroid cell positioning

We previously found that eurydendroid cell positioning is disrupted in *reelin* mutants, in which increased numbers of eurydendroid cells fail to migrate toward the Purkinje cell layer and instead remain within the granule cell layer ^23^. Because the characteristic positioning of eurydendroid cells differs from that of cerebellar output neurons in other vertebrates, these findings raised the possibility that Reelin signaling contributes to the distinctive positioning of teleost eurydendroid cells. Reelin produced by granule cells accumulates in the molecular layer, but whether eurydendroid cells themselves are capable of responding to Reelin remains unclear.

To address this possibility, we examined the expression of *vldlr*, which encodes a Reelin receptor, and *dab1a*, which encodes an intracellular adaptor required for Reelin signal transduction. Both genes have previously been reported to be expressed in the developing zebrafish cerebellum, although the identities of the expressing cell types were not determined^40^. Consistent with the established role of Reelin signaling in Purkinje cell migration, both genes were expressed in RFP-positive Purkinje cells in 5-dpf *Tg(aldoca:NTR-TagRFPT)* larvae (Figure S10). We found that *vldlr* (Figures 7A–7F) and *dab1a* (Figures 7G–7L) were also expressed in many GFP-positive eurydendroid cells in 5-dpf *lhx9^GFP^* larvae. Together with the altered eurydendroid cell positioning observed in *reelin* mutants, these findings support the possibility that eurydendroid cells respond directly to Reelin, contributing to their characteristic positioning in the zebrafish cerebellum.

**Figure 7.**
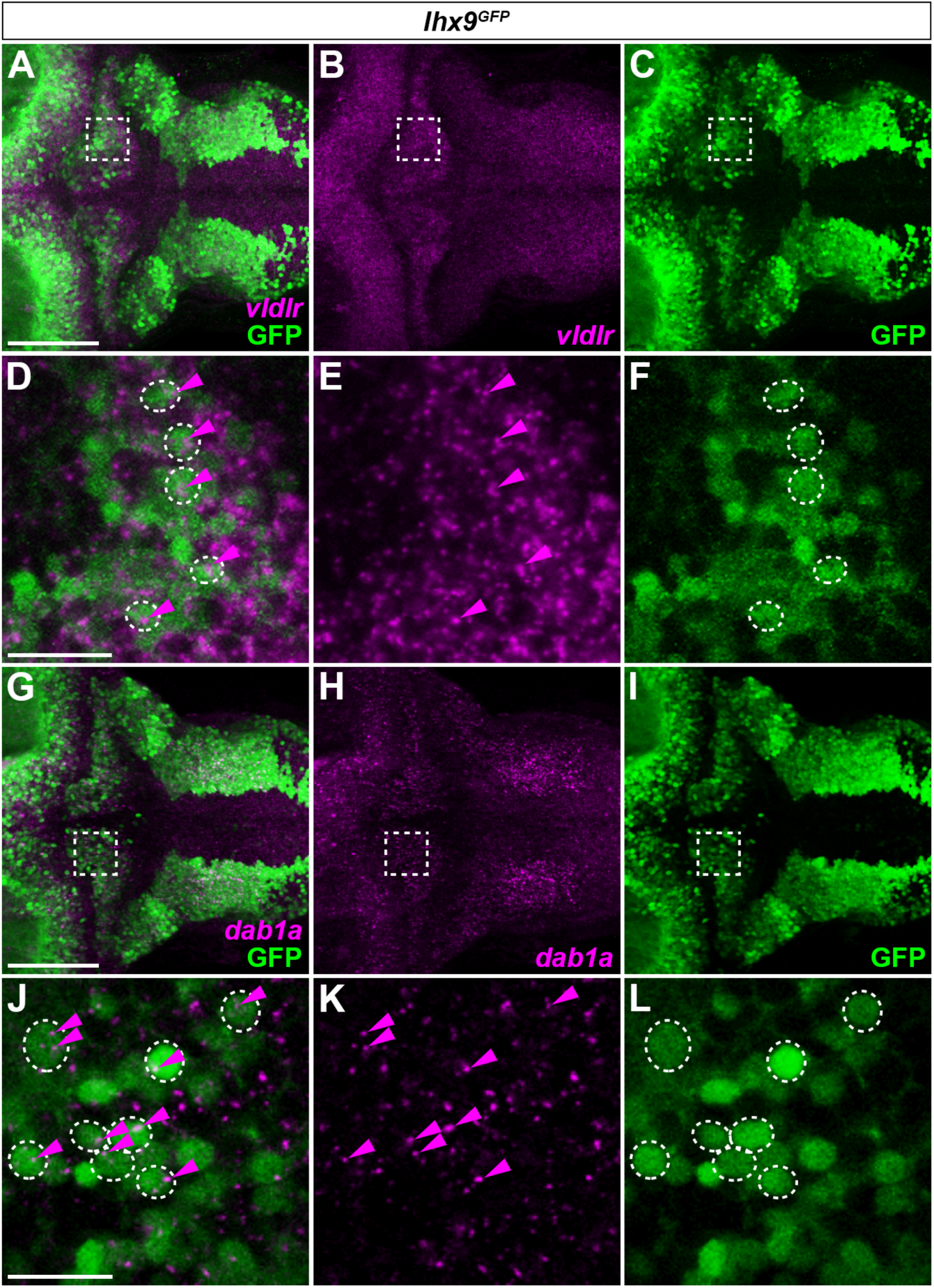
Expression of Reelin signaling molecules in eurydendroid cells. Expression of *vldlr* (magenta; A, B, D, E), *dab1a* (magenta; G, H, J, K), and GFP (green; A, C, D, F, G, I, J, L) in 5-dpf *lhx9^GFP^* larvae. Combination of hybridization chain reaction (HCR) detection of *vldlr* or *dab1a* transcripts and GFP immunostaining. Dorsal views. (D– F) Higher-magnification views of the boxed regions in (A–C). (J–L) Higher-magnification views of the boxed regions in (G–I). Dashed outlines mark GFP-positive cells expressing *vldlr* (D, F) or *dab1a* (J, L) in *lhx9^GFP^* larvae. Magenta arrowheads indicate *vldlr* or *dab1a* transcript signals within these cells. Scale bars: 50 μm (A, applies to A–C); 20 μm (D, applies to D–F); 50 μm (G, applies to G–I); 20 μm (J, applies to J–L).

## DISCUSSION

### Eurydendroid cells share molecular features with cerebellar output neurons in other vertebrates

Eurydendroid cells are considered the functional counterparts of cerebellar nuclear neurons in other vertebrates, as they receive Purkinje cell input and convey cerebellar output to extracerebellar targets. However, their organization differs markedly from that of the cerebellar nuclei. Rather than forming discrete nuclei, eurydendroid cells are distributed within the cerebellum, with their cell bodies located in the vicinity of Purkinje cells and their elaborate dendritic arbors extending into the molecular layer ^8–10^. These differences have left open the question of whether eurydendroid cells represent evolutionarily divergent cerebellar output neurons or a distinct neuronal population that acquired a cerebellar output function in the teleost lineage.

Our identification of *lhx9*, *meis2a*, and *meis2b* expression in zebrafish eurydendroid cells provides molecular evidence linking these cells to cerebellar output neurons in other vertebrates (Figures 1 and 2). *Lhx9* is expressed in cerebellar nuclear neurons in both cartilaginous fishes and mammals, although its distribution differs among vertebrate lineages; notably, *Lhx9* is absent from the cerebellar nuclei of chick and is instead expressed in extracerebellar rhombic-lip derivatives ^7,41^. *Meis2* is expressed in developing cerebellar nuclear neurons in mice and is broadly associated with the glutamatergic cerebellar nuclear lineage ^42–44^. Thus, despite their distinctive morphology and positioning, eurydendroid cells share molecular features with cerebellar output neurons across vertebrates.

The expression of these transcription factors changes during eurydendroid cell differentiation. *lhx9*, *meis2a*, and *meis2b* transcripts were initially detected broadly near the cerebellar surface, corresponding to the URL, but became progressively restricted to the anterior cerebellum by 5 dpf (Figures 1, 2, and S2). Reporter labeling persisted in adult eurydendroid cells, and *meis2a* transcripts and Meis2 protein were also detected in the adult cerebellum (Figures 1, 2, and S2). Thus, *lhx9*, *meis2a*, and *meis2b* are strongly expressed during early eurydendroid cell differentiation and persist at lower levels as these cells mature, in contrast to the more transient expression of *lhx2b* (Figure S1) ^24^.

A similar temporal organization is apparent during the development of mammalian cerebellar output neurons. Developmental single-cell transcriptomic data indicate that *Lhx2* and *Lhx9* are prominently expressed at earlier stages of the glutamatergic cerebellar nuclear lineage (Figure S12) ^26^. *Meis2* is likewise strongly expressed at early stages but, in contrast to *Lhx2* and *Lhx9*, remains broadly expressed in differentiated glutamatergic cerebellar nuclear neurons (Figure S12). These parallels suggest that, despite their markedly different anatomical organization, zebrafish eurydendroid cells and mammalian cerebellar nuclear neurons share core features of their developmental transcriptional programs.

### Eurydendroid cells share an upper rhombic lip origin with mammalian cerebellar output neurons

In mammals, glutamatergic cerebellar nuclear neurons arise from *Atoh1*-expressing progenitors in the URL, whereas GABAergic cerebellar nuclear neurons are derived from *Ptf1a*-expressing progenitors in the ventricular zone (VZ) ^14,15,45^. *Atoh1*-expressing URL progenitors generate distinct neuronal populations in a temporally regulated manner: early *Atoh1*-expressing cohorts give rise to cerebellar nuclear neurons and other early rhombic-lip derivatives, whereas later cohorts predominantly generate cerebellar granule neurons ^14^. The early-born cerebellar nuclear neurons migrate from the URL toward the anteroventral cerebellar primordium, where they contribute to the developing cerebellar nuclei (Figure 8).

**Figure 8.**
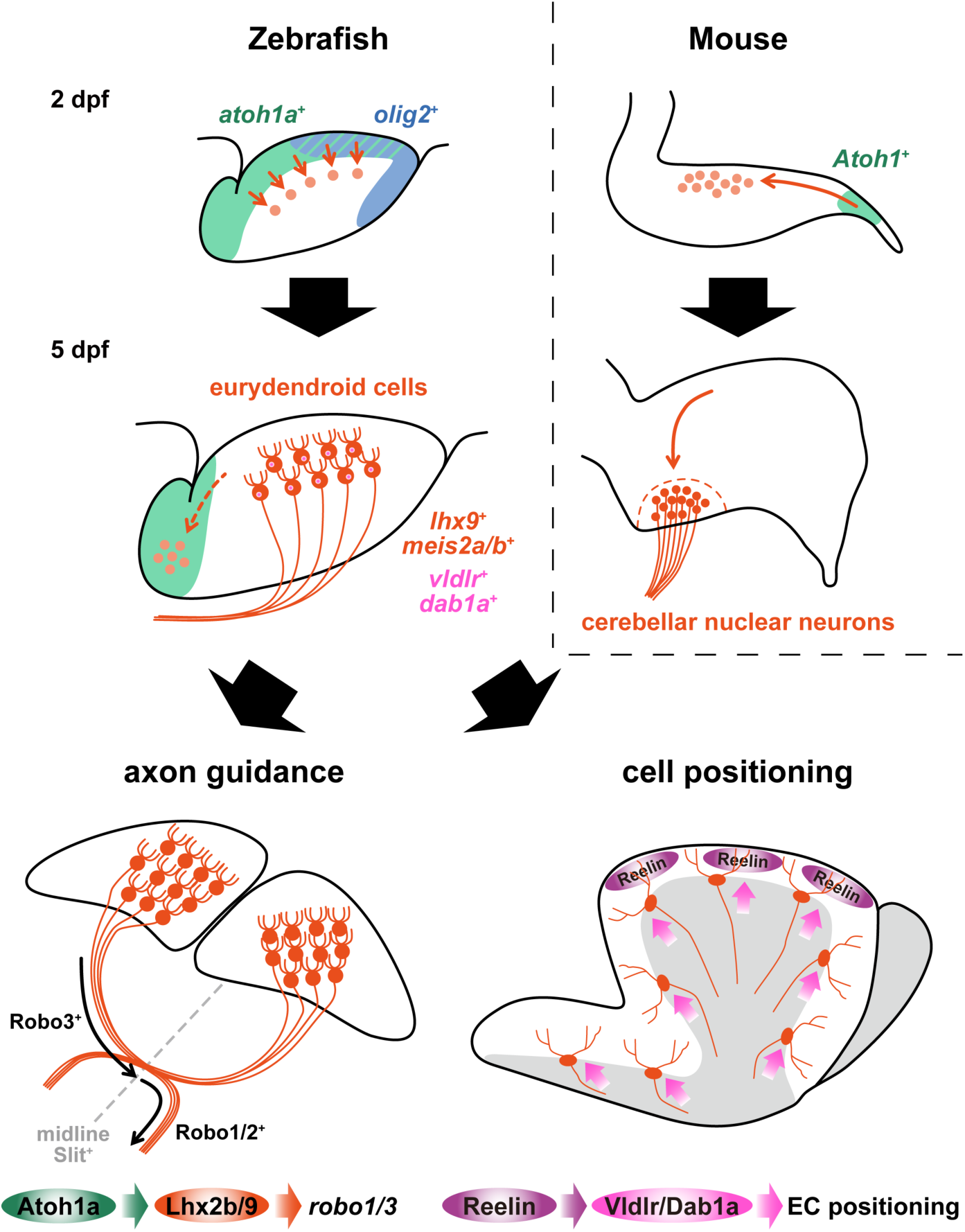
Schematic model of eurydendroid cell development. Eurydendroid cells arise from *atoh1a*-expressing upper rhombic lip progenitors, including a population expressing *olig2*. Atoh1a is required for eurydendroid cell differentiation and acts genetically upstream of *lhx2b*, *lhx9*, *meis2a*, and *meis2b*. Lhx2b and Lhx9 regulate axon guidance, at least in part through *robo1* and *robo3*, whereas Reelin signaling may contribute to the characteristic positioning of eurydendroid cells. Corresponding developmental features of mouse excitatory cerebellar nuclear neurons are shown for comparison.

Zebrafish possess three *atoh1* genes, *atoh1a*, *atoh1b*, and *atoh1c*, which exhibit distinct but partially overlapping temporal and spatial expression patterns ^19,31–33^. Among these, *atoh1a* is initially expressed broadly in the URL and subsequently becomes restricted to its anterior region (Figure 3) ^19,32,33^. This expression pattern suggests that the *atoh1a*-expressing progenitor population may resemble the early *Atoh1*-expressing progenitor population that gives rise to cerebellar nuclear neurons in mammals. Consistent with this possibility, lineage tracing revealed that many eurydendroid cells are derived from *atoh1a*-expressing progenitors (Figure 4). However, eurydendroid cells were also labeled from *atoh1b*- and *atoh1c*-expressing progenitors (Figures S3 and S4). Notably, *atoh1c*, the earliest *atoh1* gene expressed in the zebrafish cerebellar primordium, also labeled eurydendroid cells when its early-expressing population was traced (Figure S4). Thus, eurydendroid cell production is not restricted to a single *atoh1* paralog-defined lineage. Nonetheless, *atoh1a* expression delineates a URL progenitor population that gives rise to many eurydendroid cells (Figures 3 and 4). Following BrdU labeling at 5 dpf, a small number of BrdU-positive, Meis2-positive eurydendroid cells were detected at 8 dpf (Figure S11), indicating that proliferating progenitors capable of generating eurydendroid cells are still present at 5 dpf. At this stage, *atoh1a* expression is largely restricted to the anterior URL, where *lhx9* is also strongly expressed (Figures 1 and 3), raising the possibility that this anterior *atoh1a*-expressing domain continues to generate eurydendroid cells during later larval development. These findings suggest that the generation of cerebellar output neurons from *Atoh1*-expressing URL progenitors is a conserved feature of cerebellar development in teleosts and mammals.

Olig2 provides an additional link between eurydendroid cells and the developmental organization of the mammalian cerebellum. *Olig2* and *olig2* are known to be expressed in the cerebellar VZ in mammals and zebrafish, respectively ^19,21,46^. In the mammalian VZ, *Olig2* is expressed within the *Ptf1a*-positive lineage and participates in the temporal specification of GABAergic cerebellar neurons ^21^. Here, we found that *olig2* is also expressed in the zebrafish URL during early cerebellar development, where the *olig2* and *atoh1a* reporters showed overlapping labeling (Figure 3). Lineage tracing further showed that a subset of eurydendroid cells is derived from *olig2*-expressing cells, whereas very few eurydendroid cells were labeled by *ptf1a* lineage tracing (Figure S5). Together, these observations suggest that at least a subset of eurydendroid cells arises from URL progenitors that coexpress *atoh1a* and *olig2*, rather than from the *ptf1a*-expressing VZ lineage (Figure 8).

*Olig2* expression has also been associated with a subset of developing glutamatergic cerebellar nuclear neurons in mice, with a bias toward lateral cerebellar nuclei populations ^46,47^. We did not observe an obvious spatial bias in the distribution of *olig2*-positive eurydendroid cells in zebrafish (Figure 4) ^8,19^. Previous studies have also suggested molecular heterogeneity among zebrafish eurydendroid cells, identifying *olig2*-positive and calretinin-immunoreactive populations ^18,20^. Together with the previously reported molecular heterogeneity, the labeling of a subset of eurydendroid cells by *olig2* lineage tracing raises the possibility that *olig2* marks a molecularly distinct subpopulation of cerebellar output neurons in zebrafish. Further characterization will be required to determine whether this heterogeneity corresponds to the molecular subdivisions of cerebellar output neurons observed in mammals.

### Atoh1a is required for eurydendroid cell differentiation

Consistent with the prominent expression of *atoh1a* in eurydendroid cell progenitors, eurydendroid cell markers were markedly reduced in *atoh1a* mutants. In mutants carrying an allele that removes most of the bHLH domain, marker-positive eurydendroid cells were almost completely absent in some larvae, whereas a small number remained in others (Figure 5). Interestingly, early *atoh1c* expression was increased in a subset of *atoh1a* mutants (Figure S7). Moreover, *atoh1a; atoh1c* double mutants consistently showed severe reductions in eurydendroid cell markers (Figure 5). These findings suggest that Atoh1a plays the major role in eurydendroid cell differentiation and that increased *atoh1c* expression may partially compensate for the loss of Atoh1a, potentially accounting for the variable persistence of eurydendroid cells among *atoh1a* mutants. Although a subset of eurydendroid cells arises from *atoh1b*-expressing progenitors, the mutant analyses indicate that Atoh1b is largely dispensable for eurydendroid cell differentiation, whereas Atoh1a plays a predominant role (Figures 5 and S8). Comparison of the three zebrafish *atoh1* paralogs thus allowed us to identify *atoh1a* as the principal regulator of eurydendroid cell differentiation.

Consistent with a role for Atoh1a in promoting eurydendroid cell differentiation, expression of *lhx9*, *lhx2b*, *meis2a*, and *meis2b* was markedly reduced in *atoh1a* mutants (Figure 5). A functional relationship between Atoh1 and Lhx2/9 has also been observed in other neuronal populations. In the zebrafish dorsal hindbrain, Atoh1a is required for the development of a population of Lhx2/9-positive neurons ^48^, and in the lower rhombic lip, *atoh1a*-expressing progenitors give rise to *lhx2b*-expressing neurons ^49^. Similarly, in the mouse spinal cord, *Atoh1* is required for the generation of dorsal interneuron 1 (dI1) neurons that express *Lhx2* and *Lhx9* ^50^. In the developing mouse cerebellum, *Lhx2/9* and *Meis2* are expressed in postmitotic precursors of cerebellar nuclear neurons ^43^. Furthermore, transient *ATOH1* expression in human hindbrain-derived neural stem cells induces *LHX9* and, to a lesser extent, *MEIS2* expression ^51^. Although it remains unknown whether Atoh1a directly regulates these genes, these findings, together with the shared origin of zebrafish eurydendroid cells and mammalian glutamatergic cerebellar nuclear neurons from *Atoh1*-expressing URL progenitors, suggest that an *Atoh1*-dependent transcriptional program underlying cerebellar output neuron differentiation has been conserved during vertebrate evolution (Figure 8).

### Lhx9 and Lhx2b regulate axon guidance of eurydendroid cells

We found that midline crossing of eurydendroid cell axons was markedly reduced in *lhx9*; *lhx2b* double mutants, accompanied by reduced anterior extension of their axons toward the forebrain. Expression of *robo1* and *robo3*, which encode axon guidance receptors, was also reduced in *lhx9*; *lhx2b* mutants (Figure 6). Robo3 promotes commissural axon crossing by modulating responsiveness to midline guidance cues, whereas Robo1 mediates Slit-dependent repulsion and contributes to axon guidance during and after midline crossing ^52^. Thus, reduced *robo3* expression likely contributes to the impaired contralateral projection of eurydendroid cell axons in *lhx9*; *lhx2b* mutants. The reduction in anterior axon extension could be secondary to the failure of midline crossing, but reduced *robo1* expression may also contribute to this phenotype. The expression patterns of *slit1b*, *slit2*, and *slit3* extend along the midline of the zebrafish brain ^10,53^, providing potential guidance cues for anteriorly projecting eurydendroid cell axons. Together, these findings suggest that Lhx9 and Lhx2b regulate eurydendroid cell axon guidance, at least in part, by promoting *robo3* and *robo1* expression. Changes in these guidance receptors may contribute to impaired midline crossing and reduced anterior extension of eurydendroid cell axons (Figure 8).

Regulation of Robo receptor expression by Lhx2 and Lhx9 has been demonstrated in other neuronal populations. In mouse spinal dI1 commissural neurons, Lhx2 and Lhx9 are required for *Robo3* expression and contralateral axon projection, and Lhx2 binds to a conserved regulatory element in the *Robo3* locus ^54^. Moreover, ectopic expression of *Lhx2* in the spinal cord can induce *Robo3* expression ^55^. Our findings suggest that a similar Lhx2/9– Robo3 regulatory mechanism may operate in zebrafish cerebellar output neurons.

Lhx2 also regulates *Robo* receptors in other projection neurons, although the direction of this regulation appears to be context dependent. In mouse thalamic neurons, Lhx2 represses *Robo1* and *Robo2* expression, thereby regulating the guidance and topographic organization of thalamocortical axons ^56^. In contrast, cerebellar *robo1* expression was reduced in *lhx9*; *lhx2b* mutants, suggesting that Lhx9 and Lhx2b positively regulate *robo1*, either directly or indirectly. Thus, the regulation of Robo1 by Lhx2/9 may differ among neuronal populations.

Importantly, Robo signaling also controls the contralateral projection of mammalian cerebellar output neurons. In mouse cerebellofugal axons arising from cerebellar nuclear neurons, *Robo3* is required for axons to approach the midline, whereas *Robo1* and *Robo2* are required for axons to exit the midline after crossing ^57^. Our findings therefore suggest a parallel with mammalian cerebellar nuclear neurons, raising the possibility that distinct Robo receptors also contribute to the establishment of contralateral cerebellar output pathways in zebrafish. Although regulation of Robo receptors by Lhx2/9 has not been demonstrated in mammalian cerebellar nuclear neurons, our results suggest that an Lhx2/9–Robo transcriptional program contributes to cerebellar output axon guidance across vertebrates.

### Possible role of Reelin signaling in the positioning of eurydendroid cells

Reelin signaling is well established as a regulator of Purkinje cell migration and positioning in mammals ^58,59^. Although abnormalities in cerebellar nuclei organization have also been reported in *reeler* mice ^60^, whether Reelin acts directly on cerebellar nuclear neurons remains unclear. Indeed, developmental single-cell transcriptomic data show little *Dab1* expression in glutamatergic cerebellar nuclear neurons (Figure S12) ^26^. Thus, the cerebellar nuclei abnormalities observed in *reeler* mice could, at least in part, arise secondarily from disruption of cerebellar cortical organization.

In zebrafish *reelin* mutants, eurydendroid cell positioning is also disrupted, with increased numbers of eurydendroid cells located within the granule cell layer rather than near the Purkinje cell layer ^23^. Here, we found that *vldlr* and *dab1a*, encoding a Reelin receptor and its intracellular adaptor, respectively, are expressed in eurydendroid cells as well as in Purkinje cells (Figures 7 and S10). These findings suggest that eurydendroid cells are competent to respond directly to Reelin through Vldlr-Dab1a signaling. This expression pattern differs from that in the developing mouse cerebellum, where *Dab1* shows little expression in glutamatergic cerebellar nuclear neurons (Figure S12) ^26^. This difference raises the possibility that differences in Dab1 expression and Reelin responsiveness contribute to the divergent positioning of cerebellar output neurons in zebrafish and mice.

Reelin protein accumulates in the superficial molecular layer of the zebrafish cerebellum ^23^. Purkinje cells migrate from the ventricular zone toward this Reelin-rich superficial region, whereas eurydendroid cells arise near the cerebellar surface and migrate ventrally, in the opposite direction. How Reelin regulates eurydendroid cell positioning during this migration remains to be determined. One possibility is that Reelin does not simply provide a directional cue for eurydendroid cell migration but instead contributes to their retention or final positioning near the Purkinje cell layer. Eurydendroid cells may respond to Reelin through Vldlr-Dab1a signaling and thereby remain closer to the Reelin-rich superficial region (Figure 8). Such a mechanism could contribute to the characteristic positioning of teleost cerebellar output neurons near the Purkinje cell layer rather than within discrete deep cerebellar nuclei.

### Conserved and divergent programs shape cerebellar output neuron development

Our findings support a model in which zebrafish eurydendroid cells are generated from *atoh1a*-expressing progenitors and subsequently acquire a transcriptional program characterized by the expression of *lhx9*, *meis2a*, and *meis2b* (Figure 8). Together with developmental studies in other vertebrates, these findings suggest that Atoh1-dependent differentiation and Lhx-dependent axon guidance reflect conserved developmental mechanisms contributing to cerebellar output neuron development.

At the same time, our findings point to a potentially divergent mechanism that may contribute to the distinctive organization of teleost cerebellar output neurons. In zebrafish, eurydendroid cells express components of the Reelin signaling pathway, including *vldlr* and *dab1a*, raising the possibility that differences in direct responsiveness to Reelin contribute to evolutionary variation in cerebellar output neuron positioning. Intriguingly, in the early-diverging actinopterygian *Polypterus*, cerebellar output neurons possess long dendritic processes, but their cell bodies are located away from the Purkinje cell layer and form distinct cell groups ^61^. Comparative developmental and genomic analyses across vertebrates may therefore help identify the changes that gave rise to the distinctive organization of teleost cerebellar output neurons.

More broadly, cerebellar output neurons provide a useful system for investigating how evolution modifies neuronal position and morphology while retaining core developmental programs and circuit functions.

## METHODS

### RESOURCE AVAILABILITY

#### Lead contact

Requests for further information and resources should be directed to and will be fulfilled by the lead contact, Masahiko Hibi.

#### Materials availability

The materials generated in this study are available from the lead contact upon request. Zebrafish lines used in this study are being deposited in the National BioResource Project Zebrafish (NBRP Zebrafish, Japan; https://shigen.nig.ac.jp/zebra/).

#### Data and code availability

The source data underlying the figures will be deposited in Zenodo and made publicly available upon publication of the peer-reviewed article. No original code was generated in this study.

### EXPERIMENTAL MODEL DETAILS

Wild-type zebrafish of the Oregon AB genetic background were used. For immunohistochemistry and whole-mount in situ hybridization, larvae were treated with 0.003% 1-phenyl-2-thiourea (PTU; Nacalai Tesque, 27429-22) to inhibit pigmentation. Zebrafish were maintained at 28°C under a 14-h light/10-h dark cycle. Embryos and larvae were maintained in embryo medium (EM; 5 mM NaCl, 0.17 mM KCl, 0.33 mM CaCl₂, and 0.33 mM MgSO₄) ^66^.

For clarity, abbreviated names are used for zebrafish lines throughout the main text and figures; their official allele and transgene designations are provided here. The following transgenic lines were described previously: *hspzGFFgDMC156A* (referred to as *156A*), which expresses GAL4FF, a modified GAL4 transcriptional activator also referred to as GFF, in cerebellar eurydendroid cells ^10^; *Tg(5xUAS:RFP)nkuasrfp1aTg* (referred to as *UAS:RFP*) ^62^; *TgBAC(atoh1c:GAL4FF)fh430Tg* (referred to as *atoh1c:GAL4*) ^33^; *Tg(5xUAS:EGFP)nkuasgfp1aTg* (referred to as *UAS:GFP*) ^62^; *Tg(5xUAS-hsp70l:mCherry-2A-CreERT2*)*nub99Tg* (referred to as *UAS:mCherry-CreERT2*) ^35,36^; *TgBAC(olig2:EGFP)vu12Tg* (referred to as *olig2:GFP*) ^63^; *TgBAC(ptf1a:GAL4-VP16)jh16Tg* (referred to as *ptf1a:GAL4*) ^34^; *TgBAC(slc17a6b:LOXP-DsRed-LOXP-GFP)nns14Tg* (referred to as the *vglut2a* reporter) ^37^; and *Tg(aldoca:NTR-TagRFPT)nub125Tg* ^64^. The *TgBAC(olig2:GAL4FF)* transgenic line (referred to as *olig2:GAL4*) was generated in this study.

The following knock-in reporter and mutant lines were generated in this study. Where abbreviated names are used in the main text and figures, their official ZFIN designations are given in parentheses: *lhx9^GFP^* (*lhx9^nub141Et^*), *meis2b^GAL4^* (*meis2b^nub144Et^*), *meis2a^GFP^* (*meis2a^nub143Et^*), *atoh1a^GAL4^* (*atoh1a^nub138Et^*), *atoh1b^GAL4^* (*atoh1b^nub139Et^*), *atoh1a^Δ23^* (*atoh1a^nub135^*), *atoh1a^Δ4^*(*atoh1a*^nub136^), *atoh1b^fh511^*, *atoh1b^Δ2^*(*atoh1b^nub137^*), *lhx9^Δ16^*(*lhx9^nub150^*), and *lhx2b^Δ2^*(*lhx2b^nub149^*). The previously described mutant alleles *atoh1a^fh282^*and *atoh1c^fh367^* were also used ^33^.

The animal work in this study was approved by the Nagoya University Animal Experiment Committee (approval number S250015-004) and was conducted in accordance with the Regulations on Animal Experiments at Nagoya University.

### METHOD DETAILS

#### Generation of zebrafish transgenic lines

To generate the *TgBAC(olig2:GAL4FF)* transgenic line, Tol2-mediated BAC transgenesis was performed as previously described ^67^. The zebrafish *olig2*-containing BAC, ZF-danioBAC #735, was used as previously described ^63^. The BAC was modified by bacterial homologous recombination. A cassette encoding GAL4FF, together with an SV40 polyadenylation signal and a kanamycin-resistance gene flanked by FRT sites, was inserted at the *olig2* translation start site. A Tol2 transposon cassette containing homology arms corresponding to the pIndigoBAC backbone was also introduced into the BAC. The modified BAC DNA was coinjected with Tol2 transposase mRNA into one-cell-stage zebrafish embryos to generate stable transgenic fish.

Knock-in lines were generated using the CRISPR/Cas9-mediated targeted integration method described previously ^65^. The donor constructs Mbait-hs-EGFP and Mbait-hs-GFF were used for targeted integration. Mbait-hs-EGFP contains a 23-bp sequence derived from the rat *Mc4r* gene (designated Mbait), which is efficiently cleaved by CRISPR/Cas9, followed by the *hsp70l* promoter, the EGFP coding sequence, and the bovine growth hormone polyadenylation signal. Mbait-hs-GFF is identical to Mbait-hs-EGFP except that the EGFP coding sequence is replaced with the GFF coding sequence. CRISPR/Cas9 target sites were selected within 1 kb upstream of the transcription start sites of *lhx9*, *meis2a*, *meis2b*, *atoh1a*, and *atoh1b*. The target sequences for *lhx9*, *meis2a*, *meis2b*, *atoh1a*, and *atoh1b* were 5′–GGTGATGGGCAACACGAGCT–3′, 5′–TGTGACTAAAGACATTCCCG–3′, 5′–CATTTATGCGCAAAAGCGCA–3′, 5′– CCACGCGACCAGCTTGAGAA–3′, and 5′–GCCCATATAGAGCCGACATC–3′, respectively. Two types of guide RNAs were used for targeted integration. For *lhx9* and *meis2a*, single-guide RNAs (sgRNAs) targeting the sequences described above were designed as previously described ^68^. For *meis2b*, *atoh1a*, and *atoh1b*, chemically synthesized CRISPR RNAs (crRNAs) targeting the sequences described above were used in combination with trans-activating CRISPR RNA (tracrRNA). The gene-specific crRNAs, Mbait crRNA, tracrRNA, and recombinant Cas9 protein were purchased from Integrated DNA Technologies (IDT, Coralville, IA, USA). For microinjection, approximately 1 nL of a mixture containing either 20 ng/µL gene-specific sgRNA or 5 μM gene-specific crRNA:tracrRNA duplex, together with 5 μM Mbait crRNA:tracrRNA duplex, 5 μM Cas9 protein, and 20 ng/μL donor plasmid, was injected into the blastoderm of one-cell-stage zebrafish embryos. Injected fish were raised to adulthood and crossed with wild-type fish to obtain F1 progeny. For EGFP knock-in lines, F1 embryos were screened for EGFP expression. For GFF knock-in lines, F1 fish were crossed with UAS reporter fish, and the resulting progeny were examined for UAS-dependent reporter expression. Only lines in which the reporter expression was consistent with the endogenous expression pattern of the targeted gene were established and used for subsequent analyses.

#### Generation of zebrafish mutant lines

The *atoh1a^Δ23^*, *atoh1a^Δ4^*, and *atoh1b^Δ2^* mutant lines were generated using sgRNAs designed as previously described ^68^, whereas the *lhx9^Δ16^* and *lhx2b^Δ2^* mutant lines were generated using chemically synthesized crRNAs. The *atoh1b^fh511^* mutant line was generated using sgRNAs designed as described previously ^33,69^. The target sequences for *atoh1a^Δ23^*/*atoh1a^Δ4^*, *atoh1b^Δ2^*, *lhx9^Δ16^*, and *lhx2b^Δ2^*were 5′–CCAACGTCGTGCAGAAACAG–3′, 5′– TCCAGGCCCGCGGGTCACTC–3′, 5′–CAGATGGTACACGGAGTCCC–3′, and 5′–GAACATGCCTTCAATCAGCG–3′, respectively. The two target sequences used to generate the *atoh1b^fh511^* allele were 5′–GGCTGACCCGGAGTGACCCG–3′ and 5′– GGTGCGTGCGTAATTCTCCA–3′. The resulting allele carries a 25-bp deletion at one target site and a 12-bp deletion at the other. For the *atoh1a^Δ23^*, *atoh1a^Δ4^*, *atoh1b^Δ2^*, *lhx9^Δ16^*, and *lhx2b*^Δ2^ mutant lines, sgRNAs or crRNA:tracrRNA duplexes were coinjected with recombinant Cas9 protein into one-cell-stage zebrafish embryos, as previously described ^35^. Injected fish were raised to adulthood and crossed with wild-type fish to obtain F1 progeny. Genomic DNA spanning the target sites was amplified by PCR from F1 fish. Mutations in the target regions were detected by a heteroduplex mobility assay ^70^ and confirmed by sequencing. For genotyping, the following primers were used: 5′– GGCTTAAAGGAGCTGTGGGG–3′ and 5′–CAATCCGTGCATTCTTCGCC–3′ (*atoh1a^Δ23^*/*atoh1a^Δ4^*); 5′–CCCTTCAGGACTGACTGAGC–3′ and 5′– ATGCTTTACTCCCAGGACGA–3′ (*atoh1b^Δ2^*); 5′–GGTTCTCCGTGCAGAGATGT–3′ and 5′–GTCTTTCATGCCGAAATGGT–3′ (*lhx9^Δ16^*); and 5′– TGGACACTTTCGGGAGGAGT–3′ and 5′–CTTCAGAGGCAGCTTGAGGG–3′ (*atoh1b^fh511^*). For *atoh1a^Δ23^*, *lhx9^Δ16^*, and *atoh1b^fh511^*, wild-type and mutant PCR products were distinguished by agarose gel electrophoresis. For *atoh1a^Δ4^*, genotypes were distinguished by PCR product size and heteroduplex band patterns using polyacrylamide gel electrophoresis ^70^. For *atoh1b^Δ2^*, PCR products were digested with BstEII and analyzed by agarose gel electrophoresis. For *lhx2b^Δ2^*, wild-type and homozygous mutant larvae were distinguished by derived cleaved amplified polymorphic sequence (dCAPS) analysis using primers 5′–GTTTCCGCAGAACATGCCTTCAACC–3′ and 5′– CACTTCAGACAGCGCATGTG–3′. The PCR products were digested with MspI and analyzed by polyacrylamide gel electrophoresis. Genotyping of *atoh1a^fh282^* and *atoh1c^fh367^* was performed as described previously ^33^.

#### Lineage tracing

Endoxifen treatment was performed as previously described ^35,36^. Briefly, an approximately 4 μM endoxifen solution was prepared by adding 3.3 μL of a 25 mM endoxifen stock solution (Sigma-Aldrich, SML2368) dissolved in DMSO to 20 mL of EM containing 0.003% PTU. Embryos or larvae were treated from 12 to 36 hpf or for 16 h starting at 2.5 dpf, as indicated in the figure legends, and fixed at 5 dpf for analysis. Following treatment, the larvae were washed with EM and subsequently maintained in EM containing 0.003% PTU until the desired time point. Control embryos or larvae were treated with an equivalent volume of DMSO in place of the endoxifen stock solution. For lineage-tracing experiments, larvae carrying the *vglut2a* reporter were genotyped for the GAL4 driver and UAS:mCherry-CreERT2 transgenes. For genotyping, the following primers were used: 5′– GCACATCTGACAGAAGTGG–3′ and 5′–CGATACAGTCAACTGTCTTTG–3′ for *GAL4*, and 5′–TTCCCGCAGAACCTGAAGATG–3′ and 5′–CAGCGTTTTCGTTCTGCCAA–3′ for *CreERT2*. Larvae carrying both transgenes were classified as Cre⁺, whereas those lacking one or both transgenes were classified as Cre⁻. Cre⁻ larvae were used as controls for CreERT2-dependent recombination.

#### Immunostaining

For immunostaining, anti-parvalbumin 7 (1:1,000, mouse monoclonal, ascites) ^8^, anti-Meis2 (1:700, rabbit, GeneTex, GTX117564), anti-GFP (1:1,000, rat, Nacalai Tesque, 04404-84), anti-GFP (1:1,000, rabbit, MBL, 598), anti-DsRed/RFP (1:1,000, rabbit, Clontech Laboratories, 632496), and anti-BrdU (1:500, mouse, BD Biosciences, 555627) were used. Alexa Fluor 488 goat anti-rat IgG (H+L, Thermo Fisher Scientific, A-11006), CF488A goat anti-mouse IgG (H+L, Biotium, 20018), CF488A goat anti-rabbit IgG (H+L, Biotium, 20019), CF568 goat anti-rabbit IgG (H+L, Biotium, 20103), CF405M goat anti-mouse IgG (H+L, Biotium, 20340), and CF405M goat anti-rabbit IgG (H+L, Sigma-Aldrich, SAB4600461) were used as the secondary antibodies (all diluted 1:500). Larvae were immunostained as described previously ^8^, with modifications. Larvae were fixed in 4% paraformaldehyde (PFA) in PBST [phosphate-buffered saline (PBS) containing 1% Triton X-100] for 1 h at 4°C, washed three times with PBST, and permeabilized with acetone for 12 min at −30°C. The larvae were then washed three times with PBST and twice with PBS-DT (PBS containing 1% BSA, 1% DMSO, and 1% Triton X-100) and blocked in PBS-DT containing 5% goat serum for 1 h. Samples were incubated with primary antibodies overnight at 4°C. After six washes with PBS-DT, samples were incubated with secondary antibodies overnight at 4°C and washed six times with PBS-DT. Samples used to visualize axonal projections in Figures 4 and 6 were optically cleared with SeeDB reagent as previously reported ^71,72^. Fluorescence images were acquired using an LSM700 confocal laser-scanning microscope (Zeiss).

For embryonic cryosections, embryos were fixed for 3 h at 4°C and sequentially cryoprotected in 5% and 10% sucrose in PBS for 1 h each. Samples were embedded in OCT compound and cryosectioned at 10 µm. For adult brain cryosections, adult fish heads were fixed in 4% PFA in PBS for 16 h at 4°C. After fixation, the brains were dissected and gradually cryoprotected in sucrose in PBS, followed by overnight incubation in 20% sucrose in PBS. The brains were embedded in OCT compound (Sakura Finetek) and sectioned at 14 µm using a cryostat. Immunostaining of cryosections was performed as previously described^8^.

#### Colorimetric in situ hybridization

Whole-mount in situ hybridization (WISH) was performed as previously described ^8^. DIG-labeled riboprobes for *atoh1a*, *atoh1c*, and *olig2* were prepared as previously described ^8,19^. To generate riboprobes for *lhx9*, *lhx2b*, *meis2a*, and *meis2b*, gene-specific cDNA fragments were amplified by RT-PCR from total RNA isolated from early zebrafish larvae using the following primers: 5′–GGGGATCCTGGAGGAGATGGAGAGGAGA–3′ and 5′– GGGAATTCCGGACTCTTCCTCTTTCGTG–3′ (*lhx9*); 5′–TCTGAGCTCACCTGCTTCAG–3′ and 5′–TGACGTTCTCATACGCTTCG–3′ (*lhx2b*); 5′–CCCATCCCAATGTCATGC–3′ and 5′–CGTCGTGATCTCTCCAGGAC–3′ (*meis2a*); and 5′–GGGGATCCACTTCGACGACCTCTCAGGA–3′ and 5′– GGGAATTCTACCCATGTTCATCCCCATT–3′ (*meis2b*). The PCR products were subcloned into pBluescript II SK(+) (*lhx9* and *meis2b*) or pTAC-2 (*lhx2b* and *meis2a*). Antisense DIG-labeled riboprobes were synthesized by in vitro transcription using the appropriate RNA polymerases. Larvae were hybridized with DIG-labeled riboprobes overnight at 65°C and incubated overnight at 4°C with alkaline phosphatase-conjugated anti-DIG Fab fragments (1:5,000; Roche, 11093274910). BM Purple AP substrate (Roche, 11442074001) was used for color development.

Plastic sections of stained embryos were prepared using Technovit 8100 (Heraeus Kulzer) as previously described ^19^, with modifications. Embryos were immobilized in 1% agarose, dehydrated in acetone, and embedded in Technovit 8100. The resin was polymerized, and the samples were sectioned at 10 µm using a Leica RM2125 RT rotary microtome (Leica).

In situ hybridization of adult brain sections was performed as previously described^36^, with minor modifications. Adult brain sections were prepared as described above. Sections were washed five times for 5 min each with TBST [Tris-buffered saline (TBS) prepared with diethylpyrocarbonate (DEPC)-treated water and containing 0.1% Tween-20], treated with 0.2 M HCl for 10 min, and washed three times with TBST. The sections were then acetylated for 10 min in 0.1 M Tris–HCl (pH 8.0) containing 0.5% (v/v) acetic anhydride and washed three times with TBST. Sections were pre-hybridized in hybridization buffer [50% formamide, 5× saline-sodium citrate (SSC), 0.05 mg/mL heparin, 1% Tween-20, and 0.5 mg/mL tRNA] and hybridized overnight at 65°C with DIG-labeled riboprobes. Subsequent procedures were performed as described previously ^36^. Images were acquired using an Axio Imager D1 microscope equipped with an AxioCam HRc CCD camera (Zeiss).

#### Fluorescent whole-mount in situ hybridization

Fluorescent WISH using tyramide signal amplification (TSA) was performed as previously described ^73^, with minor modifications. For *robo1* and *robo3*, riboprobes were generated from pBluescript II SK(+) containing previously described cDNA fragments ^53^. Larvae were treated with 10 µg/mL proteinase K for 45 min. For *vldlr* and *dab1a*, riboprobes were prepared as previously described ^40^. Larvae were treated with 10 µg/mL proteinase K for 50–55 min. Signals were detected using TSA Plus Cyanine 3.5 (1:100, KIKO TECH, NEL763001KT) or Alexa Fluor 488 Tyramide Reagent (1:100, Thermo Fisher Scientific, B40953) diluted in 1× Plus Amplification Diluent (Akoya Biosciences, FP1498). Following the detection of *vldlr* and *dab1a* transcripts, the samples were briefly fixed, blocked, and processed for immunostaining. Fluorescence images were acquired using an LSM700 confocal laser-scanning microscope (Zeiss).

#### Whole-mount hybridization chain reaction (HCR)

For detection of *vldlr* and *dab1a* transcripts, whole-mount HCR was performed using the ISHpalette® Short Hairpin Amplifier system (Nepa Gene) according to the manufacturer’s instructions, with minor modifications. Larvae were fixed, washed with PBS, gradually dehydrated in 25%, 50%, and 75% methanol in PBS for 5 min each, washed in 100% methanol, and stored in 100% methanol at −30°C. Fixed larvae were rehydrated through a graded methanol/PBST (PBS containing 0.1% Tween-20) series, treated with 10 µg/mL proteinase K for 80 min at 20 ℃, washed, postfixed, and washed again. The samples were then prehybridized and hybridized overnight at 37°C with S41 probe sets. The sequences of the probe oligonucleotides are listed in Table S2. Following hybridization, samples were washed in 0.5× SSCT (SSC containing 0.1% Tween-20) at 37°C and preincubated in amplification buffer. The samples were then incubated with the corresponding H1 and H2 short hairpin amplifiers: ATTO550-S41 (Nepa Gene, IPL-R-S41) for *dab1a* detection (Figures 7G–7L), or S41-DIG (Nepa Gene, IPL-D-S41CM) for *vldlr* detection (Figures 7A– 7F). DIG-labeled hairpins were detected using TSA. Samples were washed three times for 10 min each with PBST (PBS containing 0.1% Tween-20) at 37°C. Beginning with overnight blocking in 2% Blocking Reagent prepared in maleic acid buffer, all subsequent steps were performed as previously described ^73^. Following transcript detection, the samples were briefly fixed, blocked, and processed for immunostaining. Fluorescence images were acquired using an LSM700 confocal laser-scanning microscope (Zeiss).

#### BrdU incorporation

BrdU incorporation was performed as previously described ^36^, with minor modifications. At 5 dpf, larvae were treated with 1% BrdU (FUJIFILM WAKO, 023-15563) in EM containing 0.003% PTU for 8 h. After BrdU treatment, larvae were washed and maintained in EM containing 0.003% PTU until 8 dpf. At 8 dpf, larvae were fixed in 4% PFA in PBS for 1 h at 4°C, treated with 2 N HCl for 30 min, washed three times with PBST (PBS containing 0.1% Triton X-100), neutralized twice with 0.1 M Na₂B₄O₇ for 5 min each, and washed three times with PBST. Immunostaining was performed as described above.

### QUANTIFICATION AND STATISTICAL ANALYSIS

The numbers of immunopositive cells within the cerebellar region were manually counted from confocal z-stack images. Cells were examined in individual optical sections to avoid double counting of the same cell across adjacent sections. The width and rostral extent of eurydendroid cell axon bundles were measured from confocal images. Rostral extent was measured from the forebrain–midbrain boundary to the anterior ends of the axons, and bundle width was measured at the midline crossing region. All measurements were performed using ZEN software (Zeiss) or Fiji image-processing software (https://fiji.sc). One-way ANOVA followed by Tukey’s multiple-comparison test was used to compare three or more groups differing in a single experimental factor, including the analyses shown in Figures 5, 6, and S8. For the lineage-tracing experiments shown in Figures 4, S3, and S4, two-way ANOVA was performed with CreERT2 expression competence (Cre⁺ or Cre⁻) and treatment (endoxifen or DMSO) as factors. Analyses were performed separately for each driver line and treatment period. Tukey’s multiple-comparison test was used for post hoc comparisons following two-way ANOVA. Data are presented as mean ± s.e.m. Sample sizes and statistical comparisons are provided in the corresponding figure legends. All statistical analyses were performed using R version 4.6.0 (R Foundation for Statistical Computing). Graphs were generated using GraphPad Prism version 5.01 (GraphPad Software).

## ACKNOWLEDGMENTS

We thank Shin-ichi Higashijima for providing plasmids used to generate CRISPR/Cas9-mediated knock-in transgenic zebrafish, Yoshihiro Yoshihara for providing plasmids used to generate *robo1* and *robo3* riboprobes, Yu Katsuyama for providing plasmids used to generate *vldlr* and *dab1a* riboprobes, and Shin-ichi Higashijima, Koichi Kawakami, Michael Parsons, and the National BioResource Project for providing transgenic zebrafish lines. We also thank Yumiko Takayanagi and Rieko Okane for assistance with fish breeding and husbandry, and Takanori Ikenaga and the members of the Hibi laboratory for helpful discussions. This work was supported by Japan Society for the Promotion of Science (JSPS) KAKENHI grants JP23K23894 and JP25K02283 to M.H. and JP23K05845 to T.S.; Japan Science and Technology Agency (JST) CREST grant JPMJCR1753 to M.H. and JST SPRING grant JPMJSP2125 to Y.K.-F.; the Terumo Life Science Foundation; and the Uehara Memorial Foundation. K.Y. is the incumbent of the Enid Barden and Aaron J. Jade Professorial Chair in Memory of Canter John Y. Jade and is supported by Minerva Foundation (714447), ERA-Net-NEURON (Tackle-CSVD) and research grants from The M. Judith Ruth Institute for Preclinical Brain Research (P137071), the Weizmann – Nella and Leon Benoziyo Center for Neurological Diseases and the Weizmann SABRA - Yeda-Sela - WRC Program, the Estate of Emile Mimran, and The Maurice and Vivienne Wohl Biology Endowment and the Brenden-Mann Women’s Innovation Impact Fund. N.M. was supported by research grants from the Estate of Olga Klein Astrachan and the Estate of Mady Dukler.

## AUTHOR CONTRIBUTIONS

M.H. conceived the study. Y.K.-F. performed the experiments and analyzed the data. Y.K.-F., I.B., and N.M. generated resources. C.U.K., K.Y., and C.B.M. provided resources. Y.K.-F. and M.H. wrote the original draft, and Y.K.-F., T.S., and M.H. reviewed and edited the manuscript. M.H. supervised the study. Y.K.-F., T.S., K.Y., and M.H. acquired funding.

## REFERENCES

1. Butler, A.B., and Hodos, W. (2005). Comparative vertebrate neuroanatomy: evolution and adaptation, 2nd Edition (Wiley-Interscience).

2. Striedter, G.F., and Northcutt, R.G. (2020). Brains through time: a natural history of vertebrates (Oxford University Press).

3. Hibi, M., Matsuda, K., Takeuchi, M., Shimizu, T., and Murakami, Y. (2017). Evolutionary mechanisms that generate morphology and neural-circuit diversity of the cerebellum. Dev Growth Differ 59, 228–243. 10.1111/dgd.12349.

4. Nieuwenhuys, R. (1967). Comparative anatomy of the cerebellum. Prog Brain Res 25, 1–93. 10.1016/S0079-6123(08)60962-0.

5. Sultan, F., and Glickstein, M. (2007). The cerebellum: Comparative and animal studies. Cerebellum 6, 168–176. 10.1080/14734220701332486.

6. Kebschull, J.M., Casoni, F., Consalez, G.G., Goldowitz, D., Hawkes, R., Ruigrok, T.J.H., Schilling, K., Wingate, R., Wu, J., Yeung, J., and Uusisaari, M.Y. (2024). Cerebellum Lecture: the Cerebellar Nuclei-Core of the Cerebellum. Cerebellum 23, 620–677. 10.1007/s12311-022-01506-0.

7. Green, M.J., and Wingate, R.J. (2014). Developmental origins of diversity in cerebellar output nuclei. Neural Dev 9, 1. 10.1186/1749-8104-9-1.

8. Bae, Y.K., Kani, S., Shimizu, T., Tanabe, K., Nojima, H., Kimura, Y., Higashijima, S., and Hibi, M. (2009). Anatomy of zebrafish cerebellum and screen for mutations affecting its development. Dev Biol 330, 406–426. 10.1016/j.ydbio.2009.04.013.

9. Ikenaga, T., Yoshida, M., and Uematsu, K. (2005). Morphology and immunohistochemistry of efferent neurons of the goldfish corpus cerebelli. J Comp Neurol 487, 300–311. 10.1002/cne.20553.

10. Takeuchi, M., Matsuda, K., Yamaguchi, S., Asakawa, K., Miyasaka, N., Lal, P., Yoshihara, Y., Koga, A., Kawakami, K., Shimizu, T., and Hibi, M. (2015). Establishment of Gal4 transgenic zebrafish lines for analysis of development of cerebellar neural circuitry. Dev Biol 397, 1–17. 10.1016/j.ydbio.2014.09.030.

11. Matsui, H., Namikawa, K., Babaryka, A., and Koster, R.W. (2014). Functional regionalization of the teleost cerebellum analyzed in vivo. Proc Natl Acad Sci U S A 111, 11846–11851. 10.1073/pnas.1403105111.

12. Matsui, H., Namikawa, K., and Koster, R.W. (2014). Identification of the zebrafish red nucleus using Wheat Germ Agglutinin transneuronal tracing. Commun Integr Biol 7, e994383. 10.4161/19420889.2014.994383.

13. Heap, L.A., Goh, C.C., Kassahn, K.S., and Scott, E.K. (2013). Cerebellar output in zebrafish: an analysis of spatial patterns and topography in eurydendroid cell projections. Front Neural Circuits 7, 53. 10.3389/fncir.2013.00053.

14. Machold, R., and Fishell, G. (2005). Math1 is expressed in temporally discrete pools of cerebellar rhombic-lip neural progenitors. Neuron 48, 17–24. 10.1016/j.neuron.2005.08.028.

15. Wang, V.Y., Rose, M.F., and Zoghbi, H.Y. (2005). Math1 expression redefines the rhombic lip derivatives and reveals novel lineages within the brainstem and cerebellum. Neuron 48, 31–43. 10.1016/j.neuron.2005.08.024.

16. Wilson, L.J., and Wingate, R.J. (2006). Temporal identity transition in the avian cerebellar rhombic lip. Dev Biol 297, 508–521. 10.1016/j.ydbio.2006.05.028.

17. Wingate, R. (2005). Math-Map(ic)s. Neuron 48, 1–4. 10.1016/j.neuron.2005.09.012.

18. Bae, Y.K., Shimizu, T., and Hibi, M. (2005). Patterning of proneuronal and inter-proneuronal domains by hairy- and enhancer of split-related genes in zebrafish neuroectoderm. Development 132, 1375–1385. 10.1242/dev.01710.

19. Kani, S., Bae, Y.K., Shimizu, T., Tanabe, K., Satou, C., Parsons, M.J., Scott, E., Higashijima, S., and Hibi, M. (2010). Proneural gene-linked neurogenesis in zebrafish cerebellum. Dev Biol 343, 1–17. 10.1016/j.ydbio.2010.03.024.

20. McFarland, K.A., Topczewska, J.M., Weidinger, G., Dorsky, R.I., and Appel, B. (2008). Hh and Wnt signaling regulate formation of olig2+ neurons in the zebrafish cerebellum. Dev Biol 318, 162–171. 10.1016/j.ydbio.2008.03.016.

21. Seto, Y., Nakatani, T., Masuyama, N., Taya, S., Kumai, M., Minaki, Y., Hamaguchi, A., Inoue, Y.U., Inoue, T., Miyashita, S., et al. (2014). Temporal identity transition from Purkinje cell progenitors to GABAergic interneuron progenitors in the cerebellum. Nat Commun 5, 3337. 10.1038/ncomms4337.

22. Matsuda, K., Yoshida, M., Kawakami, K., Hibi, M., and Shimizu, T. (2017). Granule cells control recovery from classical conditioned fear responses in the zebrafish cerebellum. Scientific reports 7, 11865. 10.1038/s41598-017-10794-0.

23. Nimura, T., Itoh, T., Hagio, H., Hayashi, T., Di Donato, V., Takeuchi, M., Itoh, T., Inoguchi, F., Sato, Y., Yamamoto, N., et al. (2019). Role of Reelin in cell positioning in the cerebellum and the cerebellum-like structure in zebrafish. Dev Biol 455, 393–408. 10.1016/j.ydbio.2019.07.010.

24. Takeuchi, M., Yamaguchi, S., Sakakibara, Y., Hayashi, T., Matsuda, K., Hara, Y., Tanegashima, C., Shimizu, T., Kuraku, S., and Hibi, M. (2017). Gene expression profiling of granule cells and Purkinje cells in the zebrafish cerebellum. J Comp Neurol 525, 1558–1585. 10.1002/cne.24114.

25. Kebschull, J.M., Richman, E.B., Ringach, N., Friedmann, D., Albarran, E., Kolluru, S.S., Jones, R.C., Allen, W.E., Wang, Y., Cho, S.W., et al. (2020). Cerebellar nuclei evolved by repeatedly duplicating a conserved cell-type set. Science 370. 10.1126/science.abd5059.

26. Sepp, M., Leiss, K., Murat, F., Okonechnikov, K., Joshi, P., Leushkin, E., Spanig, L., Mbengue, N., Schneider, C., Schmidt, J., et al. (2024). Cellular development and evolution of the mammalian cerebellum. Nature 625, 788–796. 10.1038/s41586-023-06884-x.

27. Bertuzzi, S., Porter, F.D., Pitts, A., Kumar, M., Agulnick, A., Wassif, C., and Westphal, H. (1999). Characterization of Lhx9, a novel LIM/homeobox gene expressed by the pioneer neurons in the mouse cerebral cortex. Mech Dev 81, 193–198. 10.1016/s0925-4773(98)00233-0.

28. Retaux, S., Rogard, M., Bach, I., Failli, V., and Besson, M.J. (1999). Lhx9: a novel LIM-homeodomain gene expressed in the developing forebrain. J Neurosci 19, 783–793. 10.1523/JNEUROSCI.19-02-00783.1999.

29. Akitake, C.M., Macurak, M., Halpern, M.E., and Goll, M.G. (2011). Transgenerational analysis of transcriptional silencing in zebrafish. Dev Biol 352, 191–201. 10.1016/j.ydbio.2011.01.002.

30. Nakamura, T., Jenkins, N.A., and Copeland, N.G. (1996). Identification of a new family of Pbx-related homeobox genes. Oncogene 13, 2235–2242.

31. Adolf, B., Bellipanni, G., Huber, V., and Bally-Cuif, L. (2004). atoh1.2 and beta3.1 are two new bHLH-encoding genes expressed in selective precursor cells of the zebrafish anterior hindbrain. Gene Expr Patterns 5, 35–41. 10.1016/j.modgep.2004.06.009.

32. Chaplin, N., Tendeng, C., and Wingate, R.J. (2010). Absence of an external germinal layer in zebrafish and shark reveals a distinct, anamniote ground plan of cerebellum development. J Neurosci 30, 3048–3057. 10.1523/JNEUROSCI.6201-09.2010.

33. Kidwell, C.U., Su, C.Y., Hibi, M., and Moens, C.B. (2018). Multiple zebrafish atoh1 genes specify a diversity of neuronal types in the zebrafish cerebellum. Dev Biol 438, 44–56. 10.1016/j.ydbio.2018.03.004.

34. Parsons, M.J., Pisharath, H., Yusuff, S., Moore, J.C., Siekmann, A.F., Lawson, N., and Leach, S.D. (2009). Notch-responsive cells initiate the secondary transition in larval zebrafish pancreas. Mech Dev 126, 898–912. 10.1016/j.mod.2009.07.002.

35. Itoh, T., Uehara, M., Yura, S., Wang, J.C., Fujii, Y., Nakanishi, A., Shimizu, T., and Hibi, M. (2024). Foxp and Skor family proteins control differentiation of Purkinje cells from Ptf1a- and Neurog1-expressing progenitors in zebrafish. Development 151. 10.1242/dev.202546.

36. Wang, J.C., Shimizu, T., and Hibi, M. (2025). Transforming growth factor-beta-mediated regulation of atoh1-expressing neural progenitors is involved in the generation of cerebellar granule cells in larval and adult zebrafish. Dev Growth Differ 67, 149–164. 10.1111/dgd.70002.

37. Satou, C., Kimura, Y., and Higashijima, S. (2012). Generation of multiple classes of V0 neurons in zebrafish spinal cord: progenitor heterogeneity and temporal control of neuronal diversity. J Neurosci 32, 1771–1783. 10.1523/JNEUROSCI.5500-11.2012.

38. Ikenaga, T., Yoshida, M., and Uematsu, K. (2002). Efferent connections of the cerebellum of the goldfish, Carassius auratus. Brain Behav Evol 60, 36–51. 10.1159/000064120.

39. Takeuchi, M., Yamaguchi, S., Yonemura, S., Kakiguchi, K., Sato, Y., Higashiyama, T., Shimizu, T., and Hibi, M. (2015). Type IV Collagen Controls the Axogenesis of Cerebellar Granule Cells by Regulating Basement Membrane Integrity in Zebrafish. PLoS Genet 11, e1005587. 10.1371/journal.pgen.1005587.

40. Imai, H., Oomiya, Y., Kikkawa, S., Shoji, W., Hibi, M., Terashima, T., and Katsuyama, Y. (2012). Dynamic changes in the gene expression of zebrafish Reelin receptors during embryogenesis and hatching period. Dev Growth Differ 54, 253–263. 10.1111/j.1440-169X.2012.01327.x.

41. Pose-Mendez, S., Rodriguez-Moldes, I., Candal, E., Mazan, S., and Anadon, R. (2017). A Developmental Study of the Cerebellar Nucleus in the Catshark, a Basal Gnathostome. Brain Behav Evol 89, 1–14. 10.1159/000453654.

42. Casoni, F., Croci, L., Marroni, F., Demenego, G., Marullo, C., Cremona, O., Codazzi, F., and Consalez, G.G. (2024). A spatial-temporal map of glutamatergic neurogenesis in the murine embryonic cerebellar nuclei uncovers a high degree of cellular heterogeneity. J Anat 245, 560–571. 10.1111/joa.14107.

43. Morales, D., and Hatten, M.E. (2006). Molecular markers of neuronal progenitors in the embryonic cerebellar anlage. J Neurosci 26, 12226–12236. 10.1523/JNEUROSCI.3493-06.2006.

44. Willett, R.T., Bayin, N.S., Lee, A.S., Krishnamurthy, A., Wojcinski, A., Lao, Z., Stephen, D., Rosello-Diez, A., Dauber-Decker, K.L., Orvis, G.D., et al. (2019). Cerebellar nuclei excitatory neurons regulate developmental scaling of presynaptic Purkinje cell number and organ growth. Elife 8. 10.7554/eLife.50617.

45. Hoshino, M., Nakamura, S., Mori, K., Kawauchi, T., Terao, M., Nishimura, Y.V., Fukuda, A., Fuse, T., Matsuo, N., Sone, M., et al. (2005). Ptf1a, a bHLH transcriptional gene, defines GABAergic neuronal fates in cerebellum. Neuron 47, 201–213. 10.1016/j.neuron.2005.06.007.

46. Seto, Y., Ishiwata, S., and Hoshino, M. (2014). Characterization of Olig2 expression during cerebellar development. Gene Expr Patterns 15, 1–7. 10.1016/j.gep.2014.02.001.

47. Wizeman, J.W., Guo, Q., Wilion, E.M., and Li, J.Y. (2019). Specification of diverse cell types during early neurogenesis of the mouse cerebellum. Elife 8. 10.7554/eLife.42388.

48. Sassa, T., Aizawa, H., and Okamoto, H. (2007). Visualization of two distinct classes of neurons by gad2 and zic1 promoter/enhancer elements in the dorsal hindbrain of developing zebrafish reveals neuronal connectivity related to the auditory and lateral line systems. Dev Dyn 236, 706–718. 10.1002/dvdy.21084.

49. Belzunce, I., Belmonte-Mateos, C., and Pujades, C. (2020). The interplay of atoh1 genes in the lower rhombic lip during hindbrain morphogenesis. PLoS One 15, e0228225. 10.1371/journal.pone.0228225.

50. Bermingham, N.A., Hassan, B.A., Wang, V.Y., Fernandez, M., Banfi, S., Bellen, H.J., Fritzsch, B., and Zoghbi, H.Y. (2001). Proprioceptor pathway development is dependent on Math1. Neuron 30, 411–422. 10.1016/s0896-6273(01)00305-1.

51. Tailor, J., Kittappa, R., Leto, K., Gates, M., Borel, M., Paulsen, O., Spitzer, S., Karadottir, R.T., Rossi, F., Falk, A., and Smith, A. (2013). Stem cells expanded from the human embryonic hindbrain stably retain regional specification and high neurogenic potency. J Neurosci 33, 12407–12422. 10.1523/JNEUROSCI.0130-13.2013.

52. Comer, J.D., Alvarez, S., Butler, S.J., and Kaltschmidt, J.A. (2019). Commissural axon guidance in the developing spinal cord: from Cajal to the present day. Neural Dev 14, 9. 10.1186/s13064-019-0133-1.

53. Miyasaka, N., Sato, Y., Yeo, S.Y., Hutson, L.D., Chien, C.B., Okamoto, H., and Yoshihara, Y. (2005). Robo2 is required for establishment of a precise glomerular map in the zebrafish olfactory system. Development 132, 1283–1293. 10.1242/dev.01698.

54. Wilson, S.I., Shafer, B., Lee, K.J., and Dodd, J. (2008). A molecular program for contralateral trajectory: Rig-1 control by LIM homeodomain transcription factors. Neuron 59, 413–424. 10.1016/j.neuron.2008.07.020.

55. Kawauchi, D., Muroyama, Y., Sato, T., and Saito, T. (2010). Expression of major guidance receptors is differentially regulated in spinal commissural neurons transfated by mammalian Barh genes. Dev Biol 344, 1026–1034. 10.1016/j.ydbio.2010.06.025.

56. Marcos-Mondejar, P., Peregrin, S., Li, J.Y., Carlsson, L., Tole, S., and Lopez-Bendito, G. (2012). The lhx2 transcription factor controls thalamocortical axonal guidance by specific regulation of robo1 and robo2 receptors. J Neurosci 32, 4372–4385. 10.1523/JNEUROSCI.5851-11.2012.

57. Tamada, A., Kumada, T., Zhu, Y., Matsumoto, T., Hatanaka, Y., Muguruma, K., Chen, Z., Tanabe, Y., Torigoe, M., Yamauchi, K., et al. (2008). Crucial roles of Robo proteins in midline crossing of cerebellofugal axons and lack of their up-regulation after midline crossing. Neural Dev 3, 29. 10.1186/1749-8104-3-29.

58. Miyata, T., Nakajima, K., Mikoshiba, K., and Ogawa, M. (1997). Regulation of Purkinje cell alignment by reelin as revealed with CR-50 antibody. J Neurosci 17, 3599–3609. 10.1523/JNEUROSCI.17-10-03599.1997.

59. Rice, D.S., Sheldon, M., D’Arcangelo, G., Nakajima, K., Goldowitz, D., and Curran, T. (1998). Disabled-1 acts downstream of Reelin in a signaling pathway that controls laminar organization in the mammalian brain. Development 125, 3719–3729. 10.1242/dev.125.18.3719.

60. Fink, A.J., Englund, C., Daza, R.A., Pham, D., Lau, C., Nivison, M., Kowalczyk, T., and Hevner, R.F. (2006). Development of the deep cerebellar nuclei: transcription factors and cell migration from the rhombic lip. J Neurosci 26, 3066–3076. 10.1523/JNEUROSCI.5203-05.2006.

61. Ikenaga, T., Shimomai, R., Hagio, H., Kimura, S., Matsumoto, K., Kato, D.I., Uesugi, K., Takeuchi, A., Yamamoto, N., and Hibi, M. (2022). Morphological analysis of the cerebellum and its efferent system in a basal actinopterygian fish, Polypterus senegalus. J Comp Neurol 530, 1231–1246. 10.1002/cne.25271.

62. Asakawa, K., Suster, M.L., Mizusawa, K., Nagayoshi, S., Kotani, T., Urasaki, A., Kishimoto, Y., Hibi, M., and Kawakami, K. (2008). Genetic dissection of neural circuits by Tol2 transposon-mediated Gal4 gene and enhancer trapping in zebrafish. Proc Natl Acad Sci U S A 105, 1255–1260. 10.1073/pnas.0704963105.

63. Shin, J., Park, H.C., Topczewska, J.M., Mawdsley, D.J., and Appel, B. (2003). Neural cell fate analysis in zebrafish using olig2 BAC transgenics. Methods Cell Sci 25, 7–14. 10.1023/B:MICS.0000006847.09037.3a.

64. Koyama, W., Hosomi, R., Matsuda, K., Kawakami, K., Hibi, M., and Shimizu, T. (2021). Involvement of Cerebellar Neural Circuits in Active Avoidance Conditioning in Zebrafish. eNeuro 8. 10.1523/ENEURO.0507-20.2021.

65. Kimura, Y., Hisano, Y., Kawahara, A., and Higashijima, S. (2014). Efficient generation of knock-in transgenic zebrafish carrying reporter/driver genes by CRISPR/Cas9-mediated genome engineering. Scientific reports 4, 6545. 10.1038/srep06545.

66. Westerfield, M. (2000). The zebrafish book: a guide for the laboratory use of zebrafish (Danio rerio), 4th Edition (University of Oregon Press).

67. Suster, M.L., Abe, G., Schouw, A., and Kawakami, K. (2011). Transposon-mediated BAC transgenesis in zebrafish. Nat Protoc 6, 1998–2021. 10.1038/nprot.2011.416.

68. Watakabe, I., Hashimoto, H., Kimura, Y., Yokoi, S., Naruse, K., and Higashijima, S.I. (2018). Highly efficient generation of knock-in transgenic medaka by CRISPR/Cas9-mediated genome engineering. Zoological Lett 4, 3. 10.1186/s40851-017-0086-3.

69. Shah, A.N., Davey, C.F., Whitebirch, A.C., Miller, A.C., and Moens, C.B. (2015). Rapid reverse genetic screening using CRISPR in zebrafish. Nat Methods 12, 535–540. 10.1038/nmeth.3360.

70. Ota, S., Hisano, Y., Muraki, M., Hoshijima, K., Dahlem, T.J., Grunwald, D.J., Okada, Y., and Kawahara, A. (2013). Efficient identification of TALEN-mediated genome modifications using heteroduplex mobility assays. Genes Cells 18, 450–458. 10.1111/gtc.12050.

71. Ke, M.T., Fujimoto, S., and Imai, T. (2013). SeeDB: a simple and morphology-preserving optical clearing agent for neuronal circuit reconstruction. Nat Neurosci 16, 1154–1161. 10.1038/nn.3447.

72. Ke, M.T., and Imai, T. (2014). Optical clearing of fixed brain samples using SeeDB. Curr Protoc Neurosci 66, Unit 2 22. 10.1002/0471142301.ns0222s66.

73. Wang, G.T., Pan, H.Y., Lang, W.H., Yu, Y.D., Hsieh, C.H., and Kuan, Y.S. (2021). Three-dimensional multi-gene expression maps reveal cell fate changes associated with laterality reversal of zebrafish habenula. J Neurosci Res 99, 1632–1645. 10.1002/jnr.24806.

