## Supplementary Information for "Conserved and divergent programs govern cerebellar output neuron development in zebrafish"

**Figure S1-S12 and Table S1 and S2**

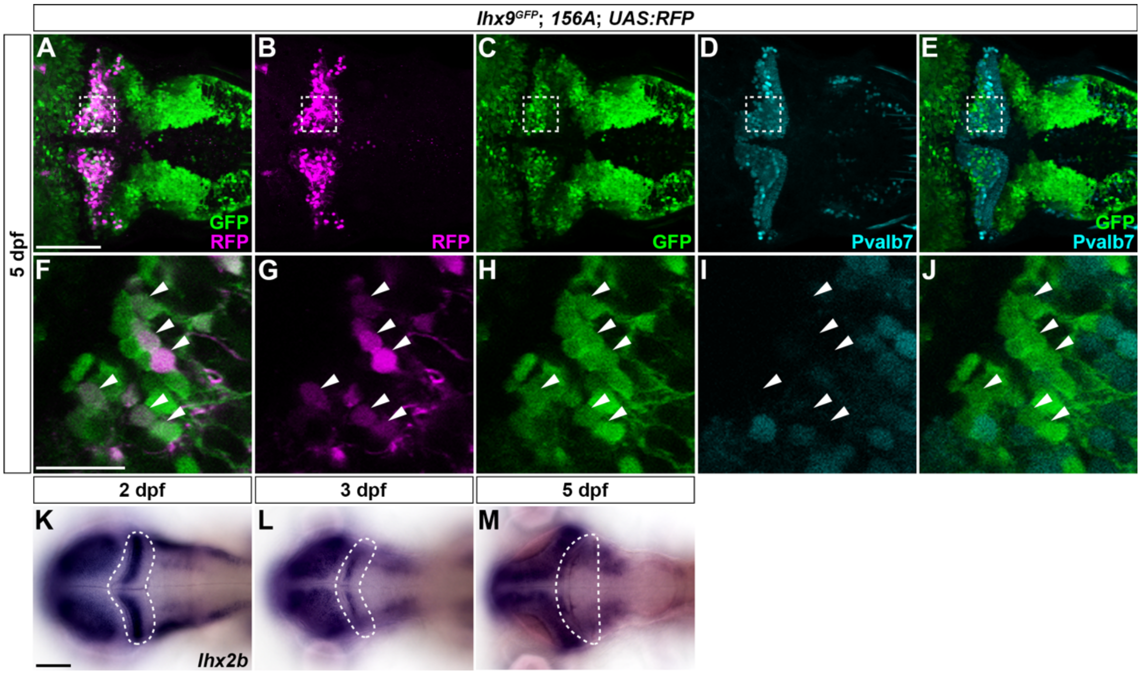


**Figure S1. Expression of *lhx9* and *lhx2b* in eurydendroid cells.**
(A–J) Expression of GFP (green; A, C, E, F, H, J), RFP (magenta; A, B, F, G), and Pvalb7 (cyan; D, E, I, J) in the cerebellum of 5-dpf *lhx9*$^{GFP}$; *hspzGFFgDMC156A*; *Tg(UAS:RFP)* larvae. Immunostaining. Dorsal views. (F–J) Higher-magnification views of the boxed regions in (A–E). Arrowheads indicate GFP- and RFP-positive but Pvalb7-negative cells. (K–M) Expression of *lhx2b* at 2 (K), 3 (L), and 5 dpf (M). In situ hybridization. Dorsal views. The cerebellar region in K–M is outlined by a dashed line. Scale bars: 50 µm (A, applies to A–E); 20 µm (F, applies to F–J); 100 µm (K, applies to K–M).

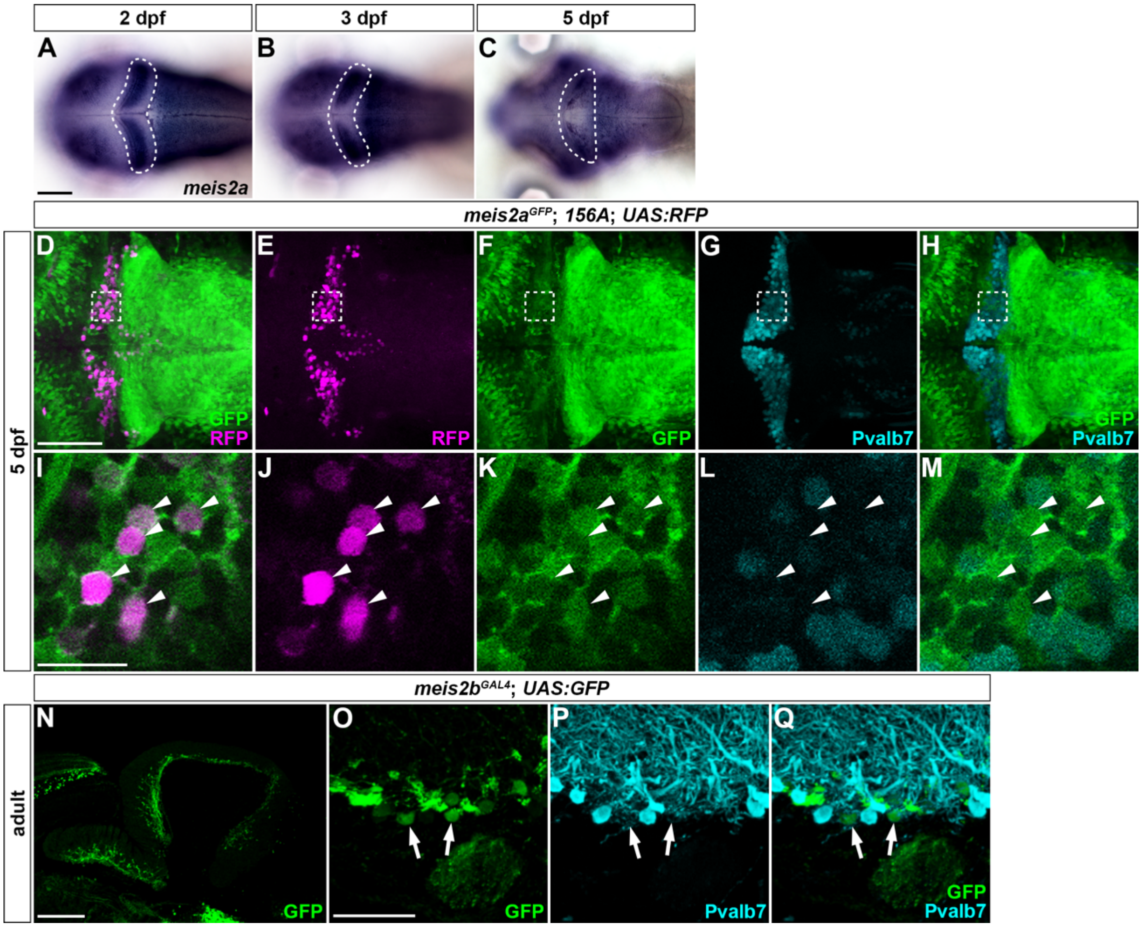


**Figure S2. Expression of *meis2a* and *meis2b* in eurydendroid cells.**
(A–C) Expression of *meis2a* at 2 (A), 3 (B), and 5 dpf (C). In situ hybridization. Dorsal views. The cerebellar region in each panel is outlined by a dashed line. (D–M) Expression of GFP (green; D, F, H, I, K, M), RFP (magenta; D, E, I, J), and Pvalb7 (cyan; G, H, L, M) in the cerebellum of 5-dpf $meis2a^{GFP}$; *hspzGFFgDMC156A*; *Tg(UAS:RFP)* larvae. Immunostaining. Dorsal views. (I–M) Higher-magnification views of the boxed regions in (D–H). Arrowheads indicate cells positive for both GFP and RFP but negative for Pvalb7. (N–Q) Expression of GFP (green; N, O, Q) and Pvalb7 (cyan; P, Q) in the cerebellum of adult $meis2b^{GAL4}$; *Tg(UAS:GFP)* zebrafish. Immunostaining of sagittal sections. (O–Q) Higher-magnification views of the region containing eurydendroid cells and the Purkinje cell layer. Arrows indicate GFP-positive cells surrounded by Pvalb7-positive axons. Scale bars: 100 µm (A, applies to A–C); 50 µm (D, applies to D–H); 20 µm (I, applies to I–M); 200 µm (N); 50 µm (O, applies to O–Q).

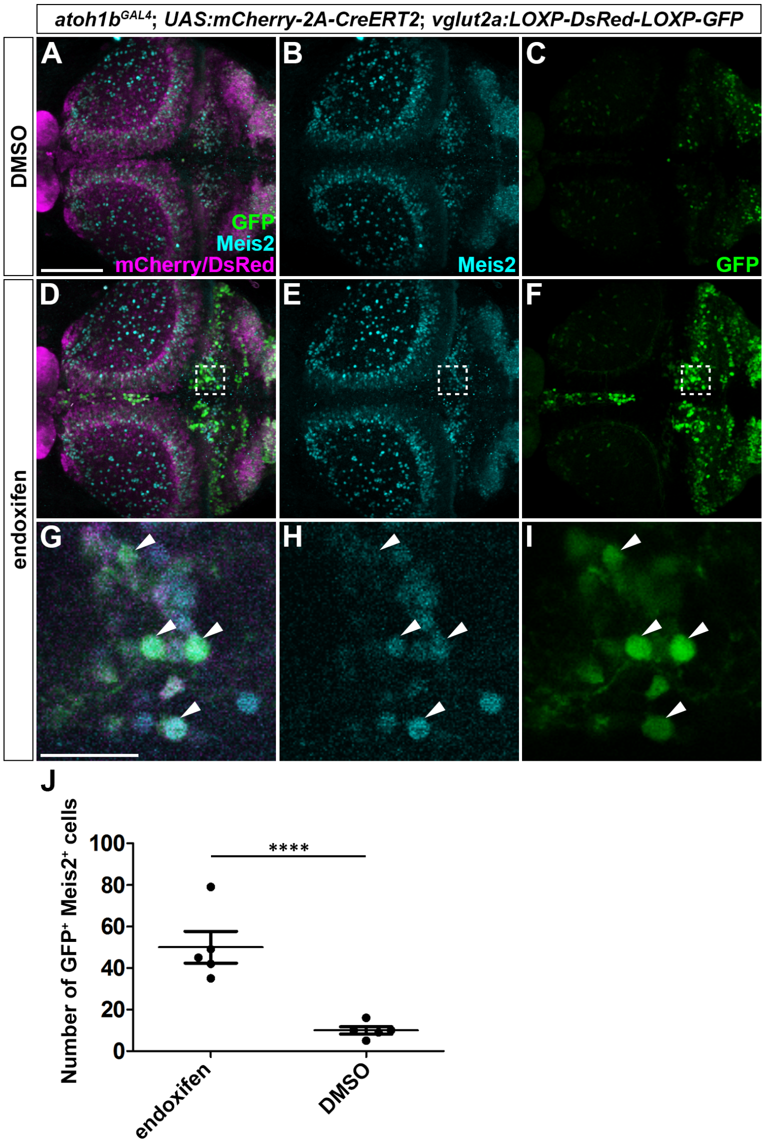


**Figure S3. Lineage tracing of *atoh1b*-expressing progenitors.**
(A–I) Expression of GFP (green), Meis2 (cyan), and mCherry/DsRed (magenta) in the cerebellar region of 5-dpf *atoh1bGAL4*; *Tg(UAS:mCherry-2A-CreERT2)*; *Tg(vglut2a:LOXP-DsRed-LOXP-GFP)* larvae. Larvae were treated with DMSO or endoxifen for 16 h from 2.5 dpf. Immunostaining for GFP and Meis2. Dorsal views. (G–I) Higher-magnification views of the boxed regions in (D–F). Arrowheads indicate cells positive for both GFP and Meis2. (J) Numbers of *vglut2a*:GFP$^{+}$ Meis2$^{+}$ cells labeled by lineage tracing. Data are shown as mean ± s.e.m., $n$ = 5 larvae per condition. Each dot represents one larva. Statistical analysis was performed using two-way ANOVA with CreERT2 expression competence (Cre$^{+}$ or Cre$^{-}$) and treatment as factors, followed by Tukey's multiple-comparison test. The analysis included the Cre$^{-}$ endoxifen- and DMSO-

treated controls shown in Figure 4, which are omitted here for clarity. ****$P < 0.0001$. Scale bars: 50 µm (A, applies to A–F); 20 µm (G, applies to G–I).

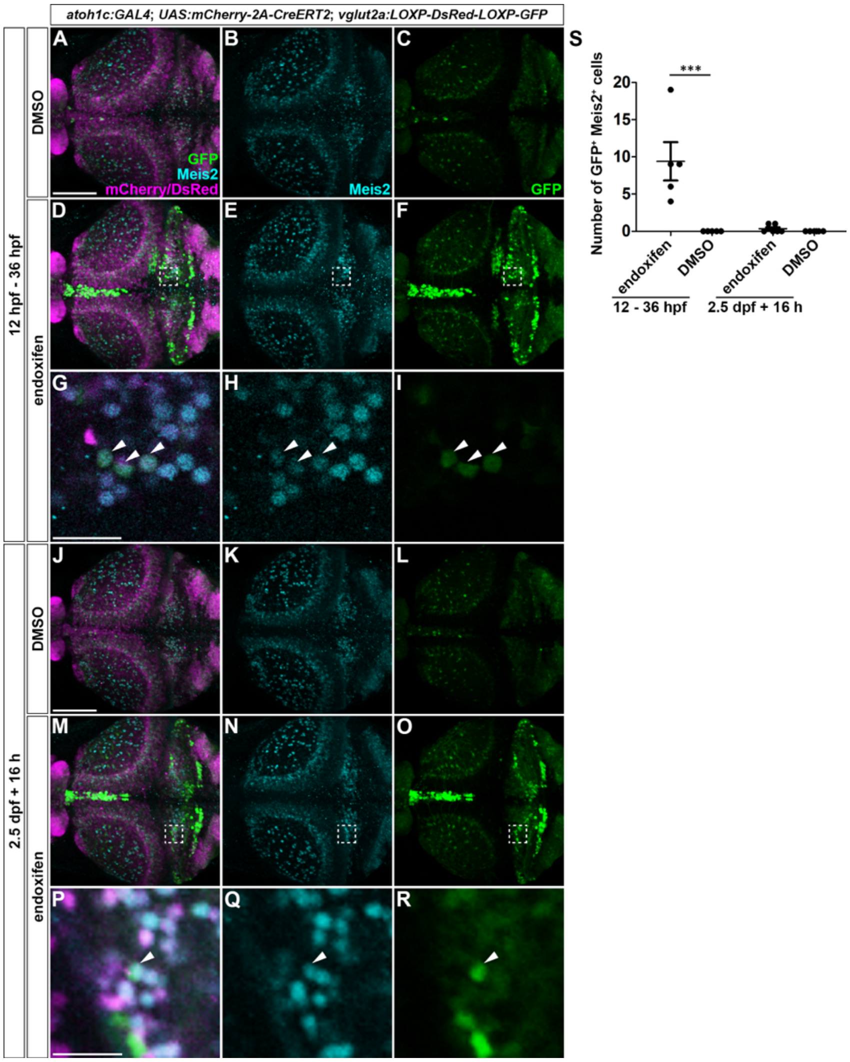


**Figure S4. Lineage tracing of *atoh1c*-expressing progenitors.**

(A–R) Expression of GFP (green), Meis2 (cyan), and mCherry/DsRed (magenta) in the cerebellar region of 5-dpf *TgBAC(atoh1c:GAL4FF)*; *Tg(UAS:mCherry-2A-CreERT2)*; *Tg(vglut2a:LOXP-DsRed-LOXP-GFP)* larvae. Larvae were treated with DMSO or endoxifen from 12 to 36 hpf (A–I) or for 16 h from 2.5 dpf (J–R). Immunostaining for GFP and Meis2. Dorsal views. (G–I) Higher-magnification views of the boxed regions in (D–F). (P–R) Higher-magnification views of the boxed regions in (M–O). Arrowheads indicate cells positive for both GFP and Meis2. (S) Numbers of *vglut2a*:GFP$^{+}$ Meis2$^{+}$ cells labeled by lineage tracing. Data are shown as mean ± s.e.m.; $n$ = 5 larvae per group for the 12–36 hpf treatment and $n$ = 6 larvae per group for the 2.5 dpf + 16 h treatment. Each dot represents one larva. Statistical analyses were performed separately for the 12–36 hpf and 2.5 dpf + 16 h treatment experiments using two-way ANOVA with CreERT2 expression competence (Cre$^{+}$ or Cre$^{-}$) and treatment as factors. Tukey's multiple-comparison test was used for post hoc comparisons following two-way ANOVA. The analyses included Cre$^{-}$

endoxifen- and DMSO-treated controls, which are not shown. The Cre$^{-}$ controls for the 2.5 dpf + 16 h treatment experiment are the same as those shown in Figure 4. ***$P < 0.001$. Scale bars: 50 μm (A, applies to A–F); 20 μm (G, applies to G–I); 50 μm (J, applies to J–O); 20 μm (P, applies to P–R).

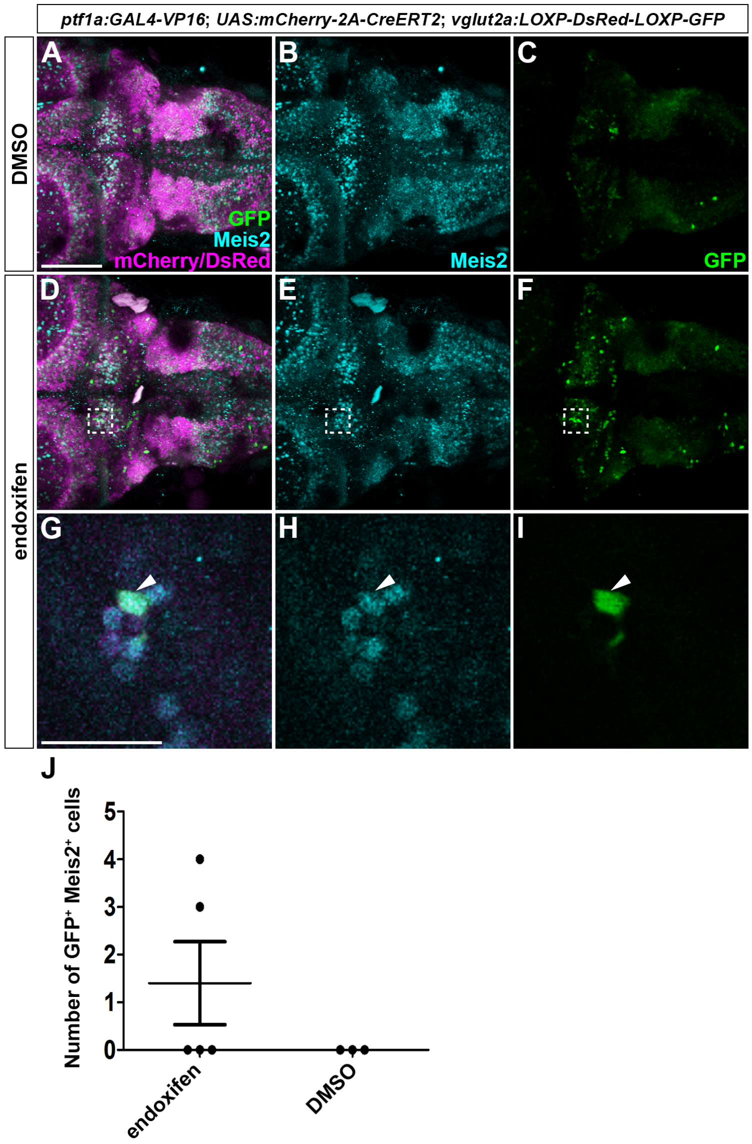


**Figure S5. Lineage tracing of *ptf1a*-expressing progenitors.**
(A–I) Expression of GFP (green), Meis2 (cyan), and mCherry/DsRed (magenta) in the cerebellar region of 5-dpf *TgBAC(ptf1a:GAL4-VP16)*; *Tg(UAS:mCherry-2A-CreERT2)*; *Tg(vglut2a:LOXP-DsRed-LOXP-GFP)* larvae. Larvae were treated with DMSO or endoxifen for 16 h from 2.5 dpf. Larvae showing GFP-positive inferior olivary neurons were selected for analysis as an indicator of effective *ptf1a*-lineage labeling. Immunostaining for GFP and Meis2. Dorsal views. (G–I) Higher-magnification views of the boxed regions in (D–F). Arrowheads indicate a cell positive for both GFP and Meis2. (J) Numbers of *vglut2a*:$GFP^{+}$ $Meis2^{+}$ cells labeled by lineage tracing. Data are shown as mean ± s.e.m.; $n$ = 5 for endoxifen-treated group and $n$ = 3 for DMSO-treated group. Each dot represents one larva. Scale bars: 50 µm (A, applies to A–F); 20 µm (G, applies to G–I).

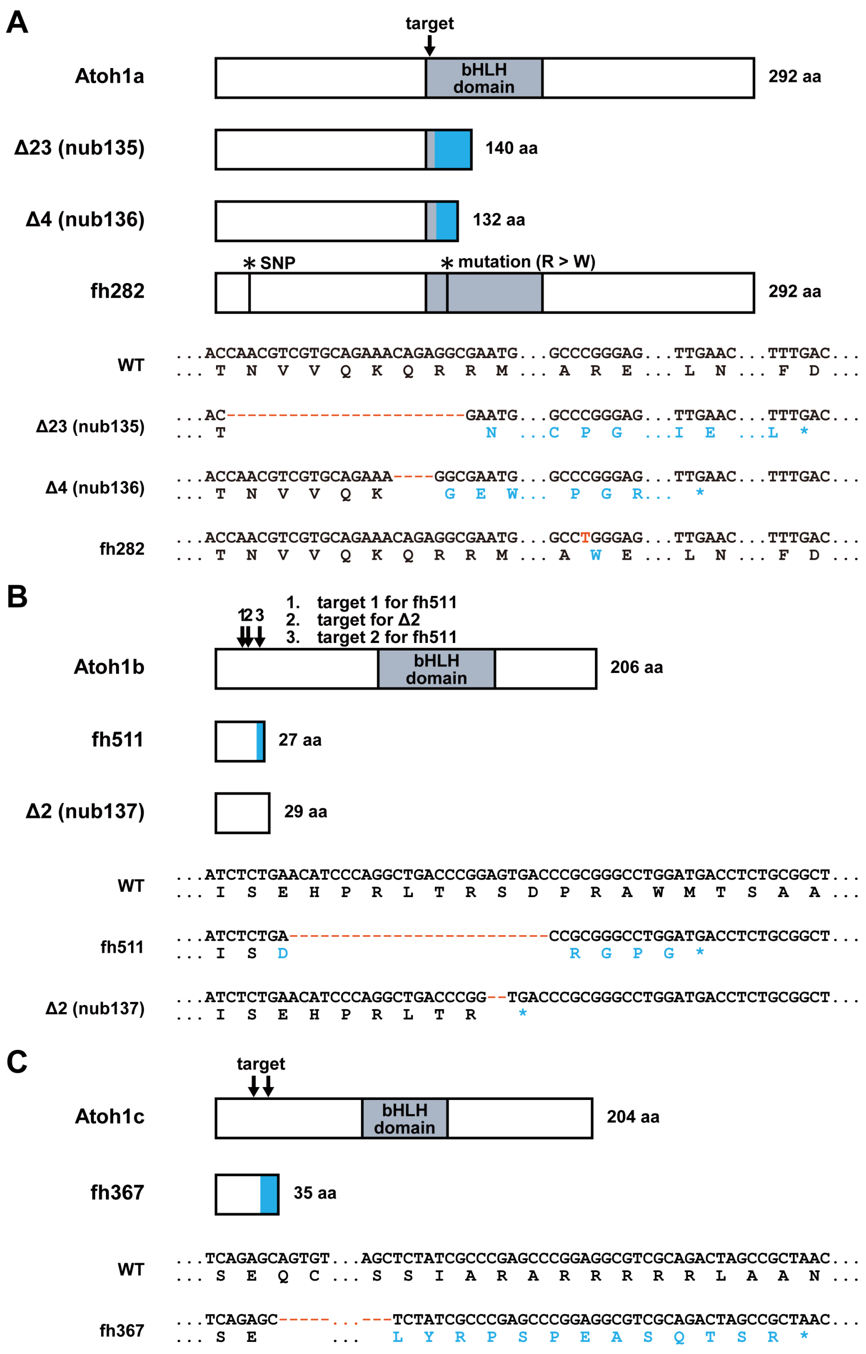


**Figure S6. Structures of wild-type and mutant Atoh1a, Atoh1b, and Atoh1c.**

Structures of wild-type (WT) and mutant Atoh1a (A), Atoh1b (B), and Atoh1c (C) proteins. The *atoh1a*$^{fh282}$ and *atoh1c*$^{fh367}$ alleles were generated by TILLING and TALENmediated mutagenesis, respectively, as described previously [33]. The other mutant alleles were generated using CRISPR/Cas9. The positions of the CRISPR/Cas9 and TALEN target sites are indicated by arrows. The *atoh1a*$^{fh282}$ allele carries an additional SNP in the region encoding the N terminus, in addition to the mutation causing an R-to-W substitution within the bHLH domain. The CRISPR/Cas9- and TALEN-generated alleles contain frameshift mutations followed by premature stop codons and are predicted to encode proteins lacking an intact bHLH domain. Altered amino acid sequences are shown in blue; blue boxes indicate frameshift-derived sequences in the protein schematics. Asterisks in the sequences indicate stop codons. Atoh1a$^{\Delta 23}$ and Atoh1a$^{\Delta 4}$ contain 28 and 15 frameshift-derived amino acids, respectively. Atoh1b$^{fh511}$ contains 5 frameshift-derived amino acids, whereas the frameshift in Atoh1b$^{\Delta 2}$ is followed immediately by a stop codon. Atoh1c$^{fh367}$ contains 13 frameshift-derived amino acids.

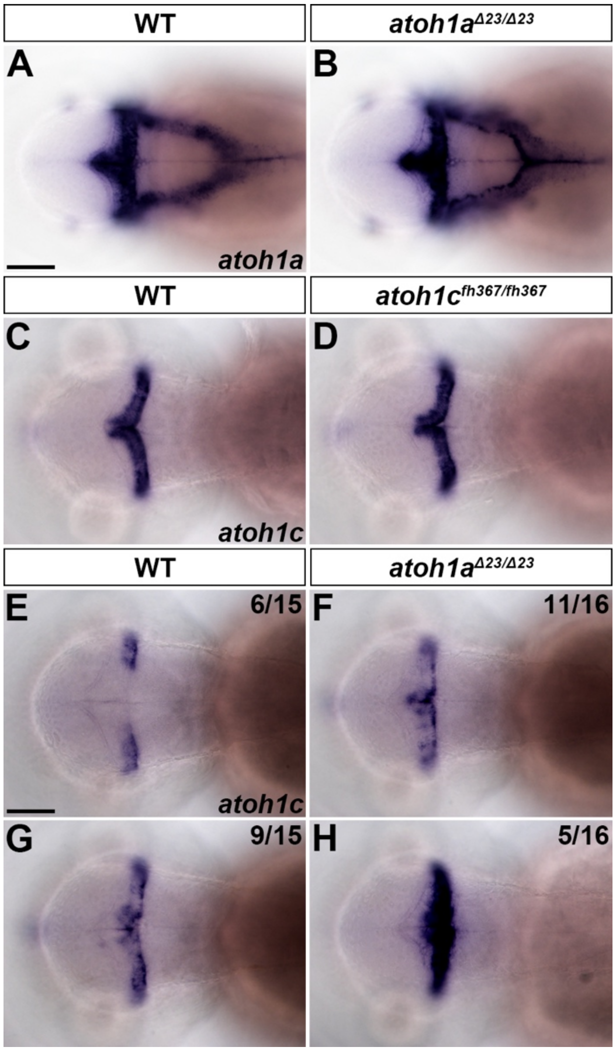


**Figure S7. Expression of *atoh1a* and *atoh1c* in *atoh1* mutants.**
(A–D) Expression of *atoh1a* in wild-type (WT; A, $n = 5$) and *atoh1a*$^{\Delta 23}$ homozygous mutant (B, $n = 5$) larvae at 2 dpf, and expression of *atoh1c* in WT (C, $n = 5$) and *atoh1c*$^{fh367}$ homozygous mutant (D, $n = 5$) larvae at 2.5 dpf. (E–H) Expression of *atoh1c* in WT (E, G) and *atoh1a*$^{\Delta 23}$ homozygous mutant (F, H) larvae at 2 dpf. *atoh1c* expression varied among WT larvae, with weak (E; 6/15 larvae) and strong (G; 9/15 larvae) expression. In *atoh1a*$^{\Delta 23}$ homozygous mutants, 11/16 larvae showed expression comparable to the strong-expression WT group (F), whereas the remaining 5/16 larvae showed markedly stronger expression (H). In situ hybridization. Dorsal views. Scale bars: 100 µm (A, applies to A–D); 100 µm (E, applies to E–H).

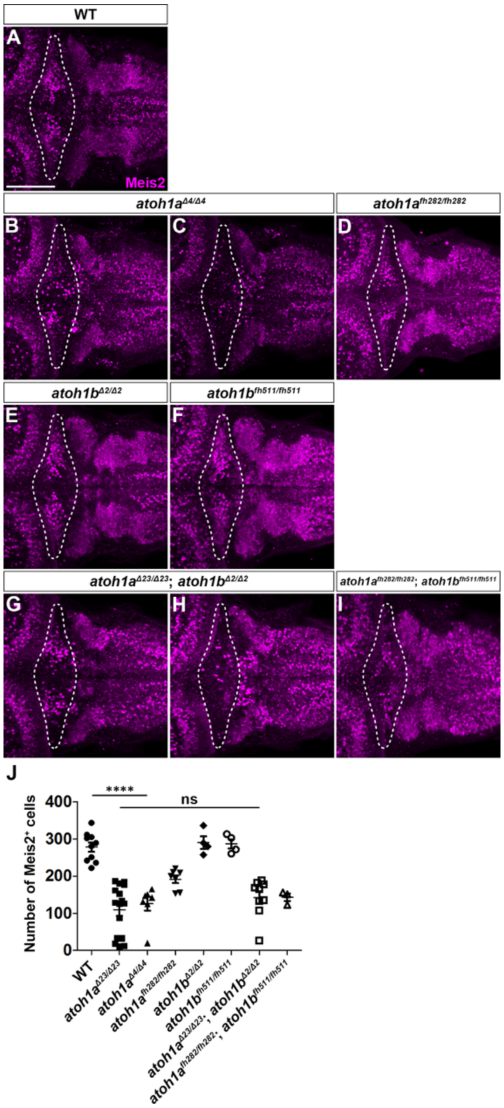


**Figure S8. Defective differentiation of Meis2-expressing eurydendroid cells in *atoh1* mutants.**

(A–I) Expression of Meis2 in 5-dpf wild-type (WT; A, *n* = 10), *atoh1a*$^{\Delta 4}$ (B, *n* = 6; C, *n* = 1), *atoh1a*$^{fh282}$ (D, *n* = 7), *atoh1b*$^{\Delta 2}$ (E, *n* = 4), *atoh1b*$^{fh511}$ (F, *n* = 4), *atoh1a*$^{\Delta 23}$; *atoh1b*$^{\Delta 2}$ (G, *n* = 8; H, *n* = 1), and *atoh1a*$^{fh282}$; *atoh1b*$^{fh511}$ (I, *n* = 3) larvae. All mutants were homozygous for the indicated alleles. Immunostaining. Dorsal views. The cerebellar region in each panel is outlined by a dashed line. Two phenotypic classes were observed in *atoh1a*$^{\Delta 4}$ and *atoh1a*$^{\Delta 23}$; *atoh1b*$^{\Delta 2}$ mutant larvae, with B and G representing larvae with more Meis2-positive cells and C and H representing larvae with fewer Meis2-positive cells, respectively. (J) Numbers of Meis2-positive cells in the cerebellum of WT and *atoh1* mutant larvae. For *atoh1a*$^{\Delta 23}$ single mutants, *n* = 15 larvae. Data are shown as mean ± s.e.m. Each symbol represents one larva. ns, not significant (*atoh1a*$^{\Delta 23}$ versus *atoh1a*$^{\Delta 23}$; *atoh1b*$^{\Delta 2}$); ****, $P < 0.0001$ (WT versus *atoh1a*$^{\Delta 4}$). One-way ANOVA with Tukey's multiple-comparison test. Scale bar: 50 µm (A, applies to A–I).

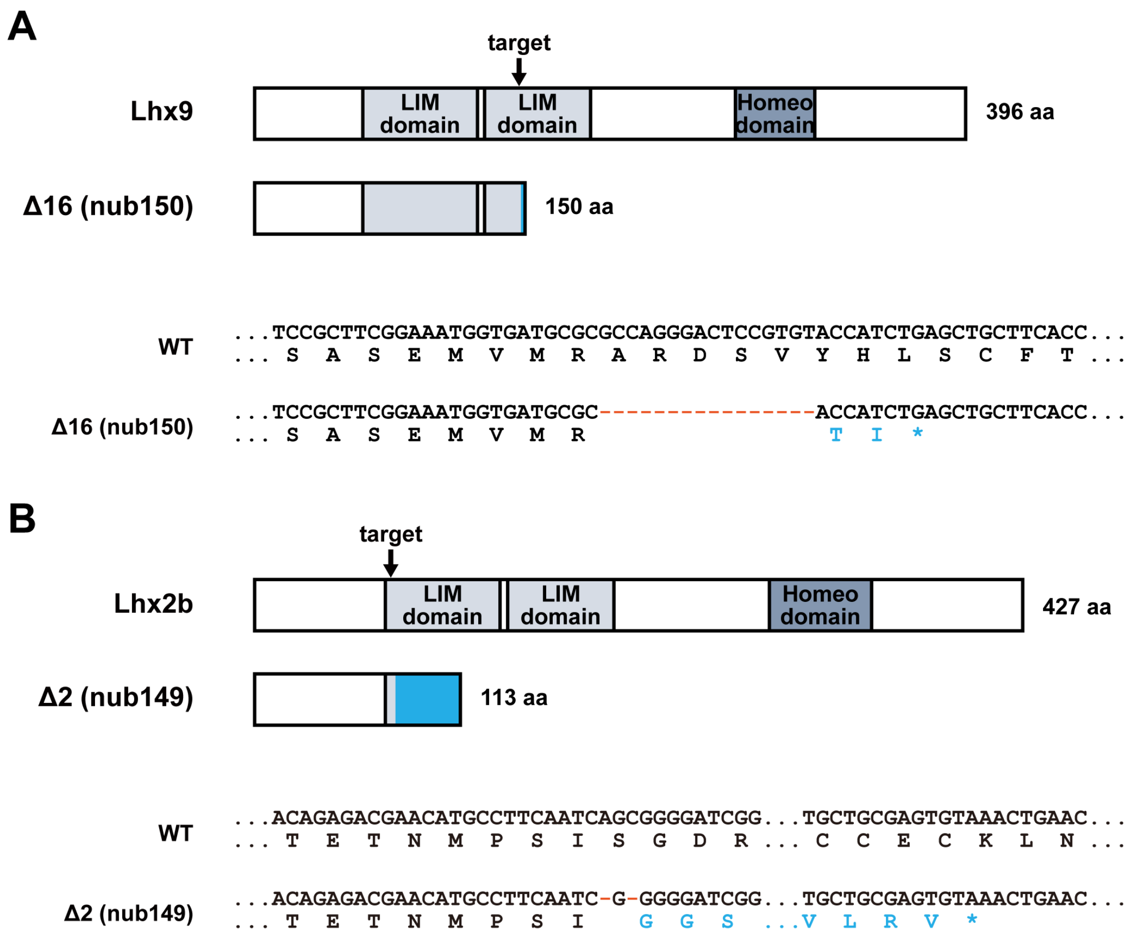


**Figure S9. Structures of wild-type and mutant Lhx9 and Lhx2b.**

Schematic protein structures and partial nucleotide and amino acid sequences of wild-type (WT) and mutant Lhx9 (A) and Lhx2b (B). The CRISPR/Cas9 target sites are indicated by arrows. The mutations cause frameshifts followed by premature stop codons. Frameshift-derived amino acid sequences are indicated by blue boxes in the schematics and blue letters in the sequences; asterisks indicate stop codons. Lhx9$^{\Delta16}$ retains the first LIM domain but lacks part of the second LIM domain and the homeodomain, whereas Lhx2b$^{\Delta2}$ lacks most of the first LIM domain, the second LIM domain, and the homeodomain. Lhx9$^{\Delta16}$ and Lhx2b$^{\Delta2}$ contain 2 and 37 frameshift-derived amino acids, respectively.

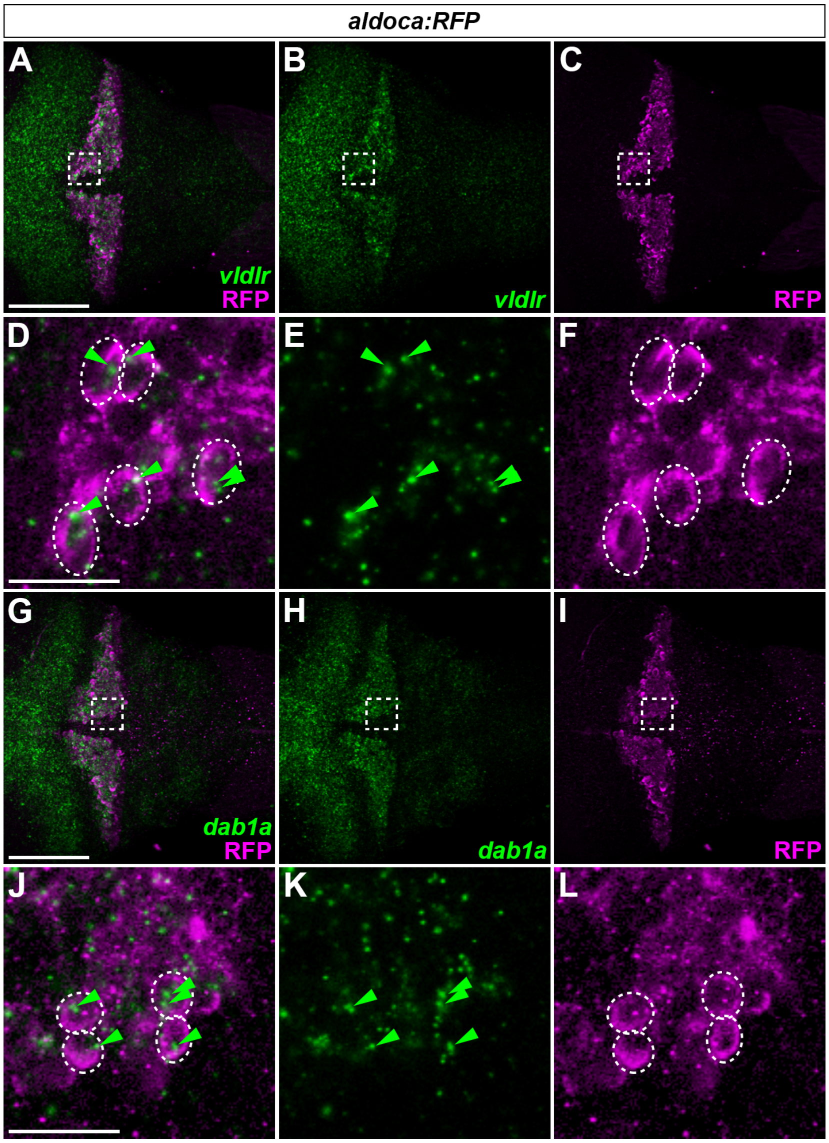


**Figure S10. Expression of *dab1a* and *vldlr* in Purkinje cells.**

Expression of *vldlr* (green; A, B, D, E), *dab1a* (green; G, H, J, K), and RFP (magenta; A, C, D, F, G, I, J, L) in the cerebellum of 5-dpf *Tg(aldoca:NTR-TagRFPT)* larvae (*aldoca:RFP* in the figure). Fluorescent whole-mount in situ hybridization for *vldlr* or *dab1a* transcripts and immunostaining for RFP. Dorsal views. (D–F) Higher-magnification views of the boxed regions in (A–C). (J–L) Higher-magnification views of the boxed regions in (G–I). Dashed outlines mark RFP-positive cells expressing *vldlr* (D, F) or *dab1a* (J, L). Green arrowheads indicate *vldlr* or *dab1a* transcript signals within these cells. Scale bars: 50 μm (A, applies to A–C); 20 μm (D, applies to D–F); 50 μm (G, applies to G–I); 20 μm (J, applies to J–L).

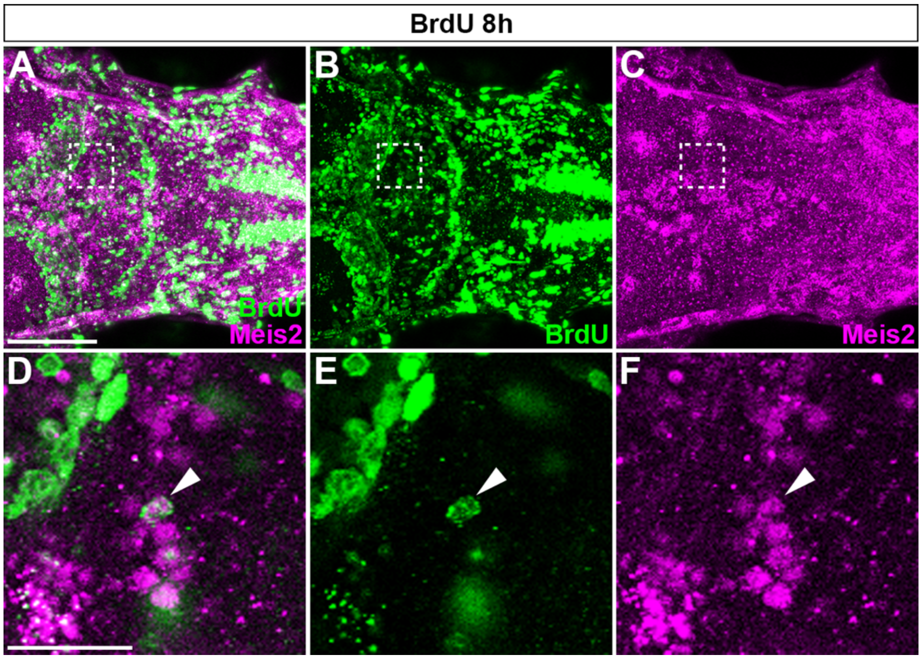


**Figure S11. Eurydendroid cells arise from cells proliferating at 5 dpf.**
(A–F) Birthdating analysis of eurydendroid cells. Larvae were treated with BrdU for 8 h at 5 dpf, fixed at 8 dpf, and immunostained with anti-BrdU (green; A, B, D, E) and anti-Meis2 (magenta; A, C, D, F) antibodies. Dorsal views. (D–F) Higher-magnification views of the boxed regions in (A–C). Arrowheads indicate a BrdU-positive, Meis2-positive cell in the cerebellum. BrdU-positive, Meis2-positive cells were detected in 2 of 8 larvae examined. Scale bars: 50 µm (A, applies to A–C); 20 µm (D, applies to D–F).

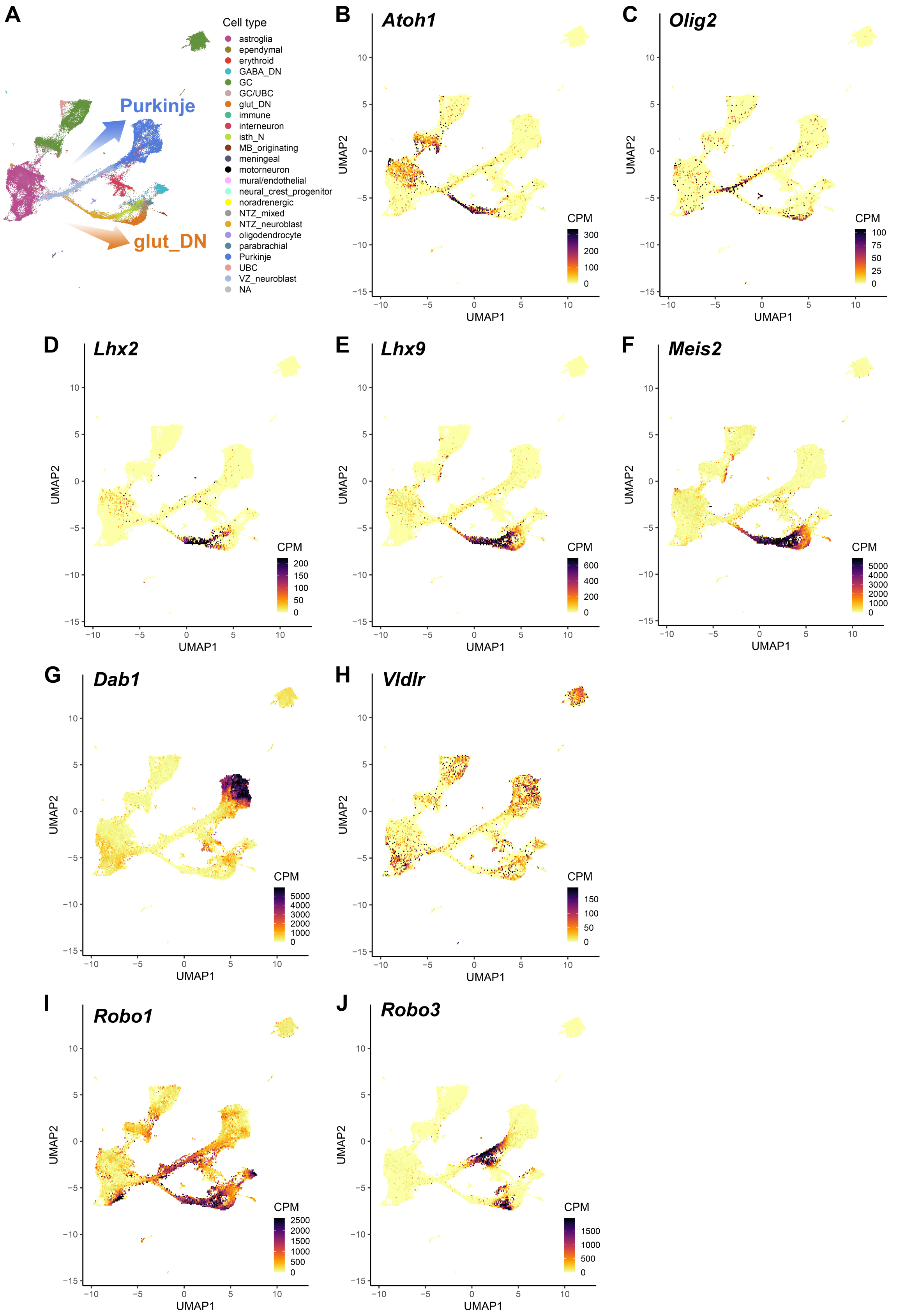

A
Cell type
astroglia
ependymal
erythroid
GABA_DN
GC
GC/UBC
glut_DN
immune
interneuron
isth_N
MB_originating
meningeal
motorneuron
mural/endothelial
neural_crest_progenitor
noradrenergic
NTZ_mixed
NTZ_neuroblast
oligodendrocyte
parabrachial
Purkinje
UBC
VZ_neuroblast
NA
Purkinje
glut_DN
B
Atoh1
C
Olig2
D
Lhx2
E
Lhx9
F
Meis2
G
Dab1
H
Vldlr
I
Robo1
J
Robo3
UMAP1
UMAP2
CPM

**Figure S12. Expression of genes associated with eurydendroid cell development in the developing mouse cerebellum.**

(A) UMAP representation of cell populations in the developing mouse cerebellum. Arrows schematically indicate the developmental progression toward glutamatergic deep cerebellar nucleus (glut_DN) neurons and Purkinje cells. (B–J) Expression of *Atoh1* (B), *Olig2* (C), *Lhx2* (D), *Lhx9* (E), *Meis2* (F), *Dab1* (G), *Vldlr* (H), *Robo1* (I), and *Robo3* (J) projected onto the UMAP. Color scales indicate expression levels in counts per million (CPM). Data were obtained from Sepp et al. [26] (https://apps.kaessmannlab.org/sc-cerebellum-transcriptome/), using the mouse dataset.

| Gene name | EC-1 | EC-2 | PC-1 | PC-2 | PC-3 | GC-1 | GC-2 |
|---|---|---|---|---|---|---|---|
| *olig2* | 101.835 | 59.4419 | 4.84757 | 1.57084 | 0.36595 | 5.64483 | 8.04205 |
| *lhx9* | 104.082 | 97.7986 | 21.2982 | 13.0048 | 13.3658 | 6.5245 | 6.27688 |
| *meis2a* | 167.95 | 84.2617 | 11.8796 | 2.78901 | 3.37331 | 5.12517 | 7.5422 |
| *meis2b* | 191.564 | 182.229 | 5.70208 | 4.48924 | 2.72466 | 3.4595 | 6.20462 |

**Table S1. Expression of selected genes in cerebellar neurons.**
Expression levels of selected genes in eurydendroid cell (EC), Purkinje cell (PC), and granule cell (GC) samples from the cell type-specific RNA-seq dataset reported previously [24]. EC (EC-1, EC-2), PC (PC-1, PC-2, and PC-3), and GC (GC-1, GC-2) samples were obtained from fluorescently labeled cells isolated from *hspzGFFgDMC156A*; *Tg(UAS:GFP)*, *Tg(aldoca:GAP-Venus)*, and *gSA2AzGFF152B*; *Tg(UAS:GFP)* transgenic larvae, respectively. EC samples may also contain neurons other than ECs labeled in *hspzGFFgDMC156A*; *Tg(UAS:GFP)* larvae. Expression levels are shown as fragments per kilobase of transcript per million mapped reads (FPKM).

| Name of oligo DNAs | Sequence |
|---|---|
| *vldlr* HCR probe | |
| *vldlr*_S41_Set1_Probe1 | gctcgacgtaaCAGGAGGAGAAGAAGCCCCAGCATA |
| *vldlr*_S41_Set1_Probe2 | AAATCCCCACAGCTGAAAACATACTaatcctttgcaaca |
| *vldlr*_S41_Set2_Probe1 | gctcgacgtaaCCGGCCATTTCCACACTGGAACTGA |
| *vldlr*_S41_Set2_Probe2 | ATCACATTGCCACACAGACGGAATaatcctttgcaaca |
| *vldlr*_S41_Set3_Probe1 | gctcgacgtaaTCCACCTCGGCACAAGTTTTCCTCA |
| *vldlr*_S41_Set3_Probe2 | ATGCATTGTCCACTGCGACAGACAAaatcctttgcaaca |
| *vldlr*_S41_Set4_Probe1 | gctcgacgtaaCACATTTCGATGCTTTCGTCTGAGC |
| *vldlr*_S41_Set4_Probe2 | TCATTCACTCGACAGGTTCGCGTATaatcctttgcaaca |
| *vldlr*_S41_Set5_Probe1 | gctcgacgtaaCTTTCTCACCGTCACACTTCCAGAA |
| *vldlr*_S41_Set5_Probe2 | TAATCTCGTCCTCTCCGTTGTCGCAaatcctttgcaaca |
| *vldlr*_S41_Set6_Probe1 | gctcgacgtaaTTTACGAGAAACACAGCGGCCACTG |
| *vldlr*_S41_Set6_Probe2 | GCAGTCATCTTCACCATTGCAGACGaatcctttgcaaca |
| *vldlr*_S41_Set7_Probe1 | gctcgacgtaaTTCGCTTGGGCTGCAGGAGGAAGGA |
| *vldlr*_S41_Set7_Probe2 | AATACACGTATTGTTCCCACAAGGGaatcctttgcaaca |
| *vldlr*_S41_Set8_Probe1 | gctcgacgtaaCTGCAGTTACGGGCAGAGCAGTTTT |
| *vldlr*_S41_Set8_Probe2 | CCGTCATCACACTTGAATTGGTCAGaatcctttgcaaca |
| *vldlr*_S41_Set9_Probe1 | gctcgacgtaaCGGACCTTATTACACACTTTACTCA |
| *vldlr*_S41_Set9_Probe2 | GGTTCATCACTCCAGTCGGGACAGTaatcctttgcaaca |
| *vldlr*_S41_Set10_Probe1 | gctcgacgtaaGTGCGAACAGCCTCCGTTATTGATC |
| *vldlr*_S41_Set10_Probe2 | GAAGCCAATCGTCAAGTCTCGACAGaatcctttgcaaca |
| *vldlr*_S41_Set11_Probe1 | gctcgacgtaaTGAGGCACTCGTTGACATCACCGCA |
| *vldlr*_S41_Set11_Probe2 | TACATATCTGGCTACATATCCCTGGaatcctttgcaaca |
| *vldlr*_S41_Set12_Probe1 | gctcgacgtaaGTGGGGTCCATCTGGTAACCATTGT |
| *vldlr*_S41_Set12_Probe2 | TTACCAACAGCTTTGCACACTCCAGaatcctttgcaaca |
| *vldlr*_S41_Set13_Probe1 | gctcgacgtaaTTACGGATGTCTCTTCGATTGGTGA |
| *vldlr*_S41_Set13_Probe2 | GTATATTCACGGCGCTCCAGGCCAAaatcctttgcaaca |
| *vldlr*_S41_Set14_Probe1 | gctcgacgtaaCTGGTTCCTTCAGGCCGCTGTTAAA |
| *vldlr*_S41_Set14_Probe2 | CTGATAGAGGGTCTACAGCAATAGAaatcctttgcaaca |
| *vldlr*_S41_Set15_Probe1 | gctcgacgtaaGTCTGTCATTCCCGTTCATACCAGA |
| *vldlr*_S41_Set15_Probe2 | ATTGAATGTCCGTCTCAACAAGGACaatcctttgcaaca |
| *vldlr*_S41_Set16_Probe1 | gctcgacgtaaTTTTCACCATCTGTCCAGAACACTC |
| *vldlr*_S41_Set16_Probe2 | AACTTGTTCGCCCCATAGATTGCCTaatcctttgcaaca |
| *vldlr*_S41_Set17_Probe1 | gctcgacgtaaACACGATGATATCCTGGGGTTCATT |
| *vldlr*_S41_Set17_Probe2 | TCCCAGACAGCTGAATTAGCTCATGaatcctttgcaaca |
| *vldlr*_S41_Set18_Probe1 | gctcgacgtaaAGTAGTTACCTGCCCTCCATCTTGA |
| *vldlr*_S41_Set18_Probe2 | TGCTTTACCTTTTGGGTGACTGGGGaatcctttgcaaca |
| *vldlr*_S41_Set19_Probe1 | gctcgacgtaaGCAGCAGATCCTCTGGCTGATGTAT |
| *vldlr*_S41_Set19_Probe2 | AATAACACAGGTAGAATCGCCCAGGaatcctttgcaaca |
| *vldlr*_S41_Set20_Probe1 | gctcgacgtaaTGGGTTGTCGAAGTTCATGCTCTTC |
| *vldlr*_S41_Set20_Probe2 | GTCTTCCTCTGTGGTCTTCAGATACaatcctttgcaaca |
| *dab1a* HCR probe | |
| *dab1a*_S41_Set1_Probe1 | gctcgacgtaaGCTGGAGTTCGGCCTCTGTTGACAT |
| *dab1a*_S41_Set1_Probe2 | CCTTTCTGACGCTGGGTCTGGCAGCaatcctttgcaaca |
| *dab1a*_S41_Set2_Probe1 | gctcgacgtaaTAAAACGCTTTATCAGAGCTGCTTC |
| *dab1a*_S41_Set2_Probe2 | TAGCCTTGTAACGAACGCCATCTCCaatcctttgcaaca |

| | |
|---|---|
| *dab1a*_S41_Set3_Probe1 | gctcgacgtaaTCCTGACAGAGTTTGTCTCCTCTAG |
| *dab1a*_S41_Set3_Probe2 | GCGATTCCCTTGAGTTTCATCATTGaatcctttgcaaca |
| *dab1a*_S41_Set4_Probe1 | gctcgacgtaaGTGTTCTCCTTTAGACCGCGCAGAG |
| *dab1a*_S41_Set4_Probe2 | GGACACAGTCAGAAAAACTTTCTGTaatcctttgcaaca |
| *dab1a*_S41_Set5_Probe1 | gctcgacgtaaTTCTCATCGAAGATCTTGATCCCCC |
| *dab1a*_S41_Set5_Probe2 | GCATGATGGTGCTGAAGTACCCCCGaatcctttgcaaca |
| *dab1a*_S41_Set6_Probe1 | gctcgacgtaaTTCGCGATGTAGGAAATCTCATGAA |
| *dab1a*_S41_Set6_Probe2 | CCAAAGGCTCTGTGGTCCGTGATATaatcctttgcaaca |
| *dab1a*_S41_Set7_Probe1 | gctcgacgtaaTTTGATGGCCACAAACCGATGGTTT |
| *dab1a*_S41_Set7_Probe2 | GATCACCGGCTCAGCAGACTGAGCCaatcctttgcaaca |
| *dab1a*_S41_Set8_Probe1 | gctcgacgtaaTCTTCTCGATCTCCTCCCTCTGCTT |
| *dab1a*_S41_Set8_Probe2 | GTTCGCACTGTTTGTCTTTCTGAGCaatcctttgcaaca |
| *dab1a*_S41_Set9_Probe1 | gctcgacgtaaTACTGATACACGGGATCCTCCACAT |
| *dab1a*_S41_Set9_Probe2 | GGCTCGTGTCCCGCCTCAAATACAAaatcctttgcaaca |
| *dab1a*_S41_Set10_Probe1 | gctcgacgtaaCTTTTGTTGGCTGGTAGGGACCTGA |
| *dab1a*_S41_Set10_Probe2 | TCGCTTTGGGACGTCATAAACTCCCaatcctttgcaaca |
| *dab1a*_S41_Set11_Probe1 | gctcgacgtaaCTGGAGGGGTGGACATGTCTCCAAA |
| *dab1a*_S41_Set11_Probe2 | ATGCTGGAGTCGAGGGCGATGTGATaatcctttgcaaca |
| *dab1a*_S41_Set12_Probe1 | gctcgacgtaaGCGGGGTTGAAGGGGGTGAACAGTT |
| *dab1a*_S41_Set12_Probe2 | ATCGTCACATAACCTGAGGGTACAGaatcctttgcaaca |
| *dab1a*_S41_Set13_Probe1 | gctcgacgtaaGGAGCCTGAGCAGCAAACTGTTGCT |
| *dab1a*_S41_Set13_Probe2 | ACTGGGGATTGGACGCCAAACGCTAaatcctttgcaaca |
| *dab1a*_S41_Set14_Probe1 | gctcgacgtaaCATGAAAGCGGCAGGAGAAAAGTGA |
| *dab1a*_S41_Set14_Probe2 | AGGAAGTGGTCCTACTGTCTGGGCAaatcctttgcaaca |
| *dab1a*_S41_Set15_Probe1 | gctcgacgtaaTCTGCAGTGTGCGCTCTCCATGTTG |
| *dab1a*_S41_Set15_Probe2 | ACATCTCCTTGCTCATCTTTGCCTGaatcctttgcaaca |
| *dab1a*_S41_Set16_Probe1 | gctcgacgtaaTCTGTGGTGCAGGATAGGCTAGGCT |
| *dab1a*_S41_Set16_Probe2 | ACTCTGCTGAAGTAGCTGGAGAAAGaatcctttgcaaca |
| *dab1a*_S41_Set17_Probe1 | gctcgacgtaaTCACAGTCATCCGTGTCTTGCGCCA |
| *dab1a*_S41_Set17_Probe2 | AGGTTCATCTGCGAGATGTCGAAGTaatcctttgcaaca |
| *dab1a*_S41_Set18_Probe1 | gctcgacgtaaGCCTCACCAAAAGAGTCATCGGTGG |
| *dab1a*_S41_Set18_Probe2 | TCTCCACTCCGACTAGGGCTGCCCTaatcctttgcaaca |
| *dab1a*_S41_Set19_Probe1 | gctcgacgtaaGGCTCTGCTTGAGGCTCACTCGCAC |
| *dab1a*_S41_Set19_Probe2 | TGAGGACTGTCCGTCTCTGAGCTCTaatcctttgcaaca |

**Table S2. Oligo DNAs for HCR.**

Uppercase letters indicate gene-specific sequences, whereas lowercase letters indicate the common sequences complementary to the S41 amplifier hairpins.
